# Alzheimer’s disease associated isoforms of human CD33 distinctively modulate microglial cell responses in 5XFAD mice

**DOI:** 10.1101/2023.07.04.547548

**Authors:** Ghazaleh Eskandari-Sedighi, Madeline Crichton, Sameera Zia, Erik Gomez, Chris D. St. Laurent, Leonardo M. Cortez, Zain H. Patel, Gaurav Sidhu, Susmita Sarkar, Vivian Aghanya, Valerie L. Sim, Qiumin Tan, Olivier Julien, Jason R. Plemel, Matthew S. Macauley

## Abstract

Microglia play diverse pathophysiological roles in Alzheimer’s disease (AD), with genetic susceptibility factors skewing microglial cell function to influence AD risk. CD33 is an immunomodulatory receptor associated with AD susceptibility through a single nucleotide polymorphism that modulates mRNA splicing, skewing protein expression from a long protein isoform (CD33M) to a short isoform (CD33m). Understanding how human CD33 isoforms differentially impact microglial cell function *in vivo* has been challenging due to functional divergence of CD33 between mice and humans. We address this challenge by studying transgenic mice expressing either of the human CD33 isoforms crossed with the 5XFAD mouse model of amyloidosis and find that human CD33 isoforms have opposing effects on the response of microglia to amyloid-β (Aβ) deposition. Mice expressing CD33M have increased Aβ levels, mo7re diffuse plaques, fewer disease-associated microglia, and more dystrophic neurites compared to control 5XFAD mice. Conversely, CD33m promotes plaque compaction and microglia-plaque contacts, and minimizes neuritic plaque pathology, highlighting an AD protective role for this isoform. Protective phenotypes driven by CD33m are detected at an earlier timepoint compared to the more aggressive pathology in CD33M mice that appears at a later timepoint, suggesting that CD33m has a more prominent impact on microglia cell function at earlier stages of disease progression. In addition to divergent roles in modulating phagocytosis, scRNAseq and proteomics analyses demonstrate that CD33m^+^ microglia upregulate nestin, an intermediate filament involved in cell migration, at plaque contact sites. Overall, our work provides new functional insights into how CD33, as a top genetic susceptibility factor for AD, modulates microglial cell function.

## Background

Late onset Alzheimer’s disease (LOAD) arises from a complex interplay between genetic and environmental factors. Genome-wide association studies (GWAS) suggest that approximately 70% of Alzheimer’s disease (AD) risk is attributable to genetic factors [1, 2]. Elucidating the functional aspects of identified genetic factors can facilitate our understanding of the underlying mechanisms contributing to disease pathogenesis. Human genetics, along with mouse models, cumulatively point to immune cells in the brain, called microglia, as being critical in AD pathogenesis through numerous functions, including their ability to phagocytose amyloid beta (Aβ) [3–7]. Of the numerous genes identified by GWAS that are linked to AD risk, a single nucleotide polymorphism (SNP) in the *CD33* gene is one of the top ranked susceptibility loci [8, 9]. In fact, among the top ten AD risk genes conferring small effects, *CD33* was the only one that showed significant effects in combination with *APOE ε4* allele, suggesting that the impact of CD33 on AD susceptibility can be more profound when combined with other major risk factors [10]. CD33 is a member of the immunomodulatory sialic acid-binding immunoglobulin-type lectin (Siglec) family that is expressed on myeloid cells. In the brain, CD33 is predominantly expressed by microglia and its immunoregulatory roles are of great interest in the context of AD pathogenesis [11, 12].

The common AD-associated SNP in *CD33,* rs3865444, is located in the promoter region and is believed to be a functional proxy of a nearby SNP, rs12459419, located four nucleotides into exon-2. This latter SNP modulates mRNA splicing [11, 12]. The common *CD33* allele (rs12459419C) favors the production of a long protein isoform known as human CD33M (CD33M; M=major) and is associated with increased susceptibility to AD relative to a minor *CD33* allele (rs12459419T) [13]. The minor AD protective *CD33* SNP enhances exon-2 skipping, which results in enhanced production of a short protein isoform known as CD33m (m=minor) that lacks the glycan binding IgV domain [8]. CD33m was originally thought to deliver its protective effects through a loss-of-function stemming from decreased expression of CD33M, however, several recent studies implicated a gain-of-function role for CD33m that may contribute to decreased AD risk. First, another variant of *CD33* (rs201074739), which is a null allele, is not AD protective [14]. Second, our group and others independently provided evidence supporting a gain-of-function role for CD33m in microglia *in vitro*, with the most striking phenotype being an enhancement in phagocytosis [15, 16]. It is noteworthy to mention that the protective *CD33* allele appears to be derived in humans and is absent in mice and non-human primates [17]. In this regard, an important consideration in studying the impact of CD33 in AD is the lack of functional conservation between mouse CD33 (mCD33) and human CD33 (hCD33) [18].

Directly testing a gain-of-function role for CD33m can be challenging as human cells express both CD33 isoforms, making it difficult to deconvolute isoform-specific phenotypes. To tackle these challenges, and to enable the study of individual hCD33 isoforms *in vivo*, we previously generated and reported two transgenic (Tg) mouse models that express either human CD33M or CD33m in the microglial cell lineage and demonstrated that the two hCD33 isoforms have opposing roles in regulating phagocytosis in microglia *in vitro* [15, 19]. Moreover, we provided evidence towards the gain-of-function effects of CD33m at functional (enhanced phagocytosis) and transcriptional levels (scRNAseq) [15]. Although the phenotypic aspects of hCD33 isoforms, such as their impact on phagocytosis have been partially explored *in vitro* [15, 16, 19], the impact of each isoform on microglial cell function *in vivo* – particularly in the context of AD pathogenesis – has not yet been investigated.

Herein, motivated to deconvolute the function of the two AD-related protein isoforms of hCD33 *in vivo*, we crossed CD33M and CD33m transgenic mice with the 5XFAD model of amyloidosis [20]. Immunofluorescence microscopy (IF), transcriptomics, proteomics, and biochemical analyses were used to examine the impact of each isoform on microglial cell response and Aβ-induced pathogenesis. We show that hCD33 isoforms differentially modulate microglia and have opposing effects on Aβ accumulation, and plaque composition. CD33M-expressing mice have more diffuse Aβ plaques while CD33m-expressing mice have a more compact plaque phenotype and show increased number of plaque-associated microglia. Accordingly, the gain-of-function role for CD33m goes beyond decreasing the amount of toxic Aβ aggregates, to increasing the number of Aβ deposits with inert dense cores, which strongly correlates with decreased number of dystrophic neurites. Overall, this study is a major step forward towards elucidating the *in vivo* roles of human-specific CD33 protein isoforms in microglia, further deciphering the functional aspects of this GWAS-identified risk factor, and helping to define early events that contribute to AD pathogenesis.

## Methods

### Animals

The Rosa26-Stopfl/fl-hCD33M/m mice were generated on a C57BL/6 genetic background and described previously [15, 19, 21]. The CX3CR1Cre (B6J.B6N(Cg)- Cx3cr1tm1.1(cre)Jung/J) and 5XFAD (B6.Cg-Tg(APPSwFlLon,PSEN1*M146L*L286V) 6799Vas/Mmjax) mice were obtained from the Jackson Laboratory. To generate 5XFAD mice expressing either hCD33M (Cx3cr1Cre+/+-hCD33M+/−) or hCD33m (Cx3cr1Cre+/+-hCD33m+/−), we crossed (Cx3cr1Cre+/+-hCD33M-/+ or Cx3cr1Cre+/+-hCD33m-/+ mice with 5XFAD-/+ mice and confirmed the progeny containing hCD33 and 5XFAD with designated primers [15, 19, 21].

Breeders containing either of the hCD33 transgenes were homozygous for CX3CR1^Cre^ so that all the mice contained a single copy of CX3CR1^Cre^. All animals were maintained in ventilated racks (Tecniplast, Green Line) and cage environmental enrichment comprising 5 cm diameter plastic tubes and nesting material (“Nestlets”, Ancare Inc.). Animals were fed irradiated chow (LabDiets, 5053) and were housed with a 12 h/12 h light/dark cycle. All protocols were in accordance with the Canadian Council on Animal Care (CCAC) and were approved by the Animal Care and Use Committee at the University of Alberta.

### Immunofluorescence (IF) staining

Half brain sections were fixed in 4% PFA at 4 °C for 24 hr, followed by incubation with 30% sucrose (4 °C) for a minimum of 72 hr. The tissue was then embedded in embedding medium for frozen tissue specimens (OCT, Thermo Scientific) and stored at -80 °C until being further processed by cryostat (Thermo Scientific). Coronal sections (20 μm) within the hippocampus region were collected and a minimum of seven sections per sample were mounted onto Superfrost Plus microscope slides (Thermo Scientific) and stored at -80 °C.

For IF staining, slides were removed from -80 °C, allowed to adjust to room temperature for 20 min and washed with PBS prior to antigen retrieval. Slides were then incubated with PBS containing 5% goat serum and 0.1% Triton X-100 for 10 min at room temperature. Slides were then further treated with blocking solution containing 5% goat serum in PBS-T (PBS containing 0.2% Tween-20), followed by incubation with 500 μl of 5% goat serum containing primary antibodies overnight at 4 °C. The following primary antibodies were used in our experiments: anti-Iba-1 (rabbit monoclonal, FUJIFILM Wako Chemicals, 1:600 dilution), and anti-Aβ (MOAB-2, abcam, 1:1000), anti-Ki67 (B56, abcam, 1:50) and anti-LAMP1 (1D4B, abcam, 1:200). The slides were washed three times in PBS-T the following day and incubated with the secondary antibodies (AF568-conjugated anti-rabbit, and AF647-conjugated anti-mouse, all used at 1:500 dilution) for 1 hr, followed by three more washes in PBS-T. To minimize the fluorescent background, autofluorescence quenching kit (TrueVIEW) was used as per the manufacturer’s protocol. Lastly, the slides were incubated with Hoechst (1:2000 dilution of 10 mg/ml stock solution) for 15 min and cover-slipped with permanent mounting medium (TrueVIEW).

Thioflavin-S (Thio-S) staining of amyloid aggregates was performed as described previously [22]. Briefly, sections were stained with 150 µM Thioflavin-S (Sigma) solution in 40% ethanol for 10 min at room temperature. Slides were then washed with 50% ethanol, followed by two washes in PBS prior to mounting coverslips.

### Microscopy

Fluorescence microscopy was performed with the LSM 700 laser scanning confocal microscope (ZEISS), equipped with Axiocam 702 mono camera (ZEISS) and the images were captured at 10X magnification. A minimum of five brain sections from each animal were assessed for analysis. Confocal microscopy images were captured with the same microscope in confocal mode (software: Zen2.6 Black edition, ZEISS) and at 63X magnification oil immersion objective (N.A. 1.4) at 1024 x 1024 pixel resolution. Sample identity was blinded for all analyses and images were processed with Zen2.6 Blue edition software (ZEISS).

### Image analyses and quantification

All the global analyses were performed on widefield images captured from the hemi-brains. This included Aβ plaque burden, number of Aβ clusters, percentage of ThioS^+^ clusters, and IBA-1 density in the whole brain as well as region-specific analysis of overall Aβ levels. In this regard, the total area of the detected signal was quantified and normalized to the total area of the brain frame. For the analyses performed on individual plaques, images captured in the confocal were used. The details of analyses performed on plaques are as follows:

#### Aβ and Thioflavin-S area for individual plaques

For measuring total Aβ area within each cluster, the plaque area was selected within a frame and Aβ or ThioS fluorescent signal was distinguished by thresholding. The total area of Aβ/ThioS within the frame was then measured and recorded per plaque. Classification of Thio-S positive plaques were done by scrolling through the z-stack and judging manually. Plaque compaction was measured through dividing the Thioflavin-S area by the total Aβ area. This was calculated both globally (in high plaque density regions) as well as for individual plaque clusters.

#### Plaque-associated microglia

The area of each plaque cluster was selected by a frame and the total number of plaque-associated microglia was calculated by counting the total number of IBA-1 positive nuclei (stained with Hoechst) within the allocated frame region. To measure plaque-associated microglia density, this number was divided by the area of plaque within the frame region.

#### Microglia-plaque interface

The area of each ThioS positive plaque cluster was selected by a frame and the thresholding for the ThioS channel was adjusted in a way that only the perimeter of the core would be quantified. The IBA-1 signal overlapping with ThioS within the perimeter was then measured by the software and the overlap was quantified as a percentage of total perimeter.

#### Percentage of internalized Aβ by microglia

Microglia interacting with Aβ clusters were selected and a barrier mask for the cells was defined based on the IBA-1 signal. The total area of Aβ within the mask was measured, quantified and normalized to the total area of IBA-1.

#### Percentage of proliferative microglia (Ki67 staining)

The IBA-1^+^ cell bodies surrounding plaques (PAM) were counted and the number of Ki67^+^ cells was recorded. The proliferative population was then calculated as the percentage of total PAM that were Ki67^+^.

#### Quantification of LAMP1+ in dystrophic neurites (DNs) area in dorsal subiculum

The total area of LAMP1^+^ spheroids within the dorsal subiculum was measured and quantified as the percentage of the total brain frame.

#### Quantification of DN area in individual neuritic plaques (μm^2^)

The total area of spheroid within each neuritic plaque was measured and recorded. A total of 400 plaques from 10 mice per cohort were used for this analysis.

### Extraction of soluble and insoluble Aβ from mouse brain

Half brain samples were individually homogenized in sterile PBS containing protease (cOmplete, Roche) and phosphatase inhibitor cocktails (Thermo Fisher Scientific). Homogenization was done using ceramic magnetic beads (2.8-mm ceramic beads; Bertin Technologies SAS) in an Omni bead Ruptor system (3.2 M/s shake speed, 10s rupture, 10 s break, three repeats) to obtain 10% (w/v) homogenates.

Extraction of soluble and insoluble Aβ was done as described previously with slight modifications [23]. Briefly, 130 µL of homogenate was thawed on ice and mixed with equal amount of 2% Triton X-100, to achieve a final concentration of 1% in the homogenate. Samples were then incubated on ice for 15 min, while being vortexed every 5 min, followed by ultracentrifugation at 100,000 rcf (4 °C for 15 min). The supernatant was extracted and tested for total protein concentration with BCA assay. The pellet (insoluble fraction) was either resuspended in PBS and sonicated for 1 min for proteinase K (PK) treatment, or monomerized by addition of formic acid (final concentration of 70% (v/v). The volume of PBS used for resuspension of the pellet was adjusted based on the protein concentration of the soluble fraction.

For PK digestion, the insoluble fraction was aliquoted and treated with PK for 1 hr at 37 °C. The remaining material were then monomerized by addition of formic acid at a final concentration of 70% (v/v) and sonicated for 1 min. The solutions were neutralized (1:20) with neutralization buffer (1M of Tris base, 0.5 M of Na2HPO4, 0.05% NaN3 (w/v)) prior to measurements. Aβ levels were quantified by an electrochemiluminescence-linked immunoassay (Meso Scale Discovery (MSD), Assay 2) as per manufacturer’s protocol. The plates were read on the SECTOR Imager 6000 and data analysis was performed using the MSD DISCOVERY WORKBENCH software v.2.0.

### Proteomics

#### Tissue homogenization and sample preparation

For whole proteome analysis of each mouse genotype, individual half brains (n=5 male mice per group) were homogenized using a glass homogenizer in urea-based lysis buffer (8 M urea, 100 mM Tris pH 8.5, 1% SDS, 5 mM EDTA, 1 mM AEBSF, 1 mM PMSF and 4 mM IAM) to obtain 10% w/v homogenates. A round of probe sonication (30% amp, 2s on/ 2s off, 2 min) was applied, followed by incubation in the dark on ice for 15 min. Lysates were clarified by centrifugation and total protein concentration of each sample was determined in the recovered supernatant by the BCA assay.

A total of 50 µg of protein from each brain lysate were processed using ProTrap XG cartridges [24] (Proteoform Scientific inc.) following the manufacturer’s protocol with some modifications. Briefly, 100 mM NaCl was added into each sample and the volume was adjusted to 100 µL. Proteins were precipitated in acetone (1:4 ratio) directly on the filtration cartridge at R.T. for 30 min. The cartridge was spun down at 2500 rcf for 2 min and protein pellet was washed once with acetone. The pellet was resuspended in 100 µL of 8 M urea by vortexing for 30 s, bath sonication for 10 min, and incubation at R.T. for 30 min. The urea in the samples was diluted by addition of 400 µL of 100 mM Tris buffer (pH 8). Proteins were reduced (10 mM DTT) and alkylated (25 mM iodoacetamide) at 37°C for 30 min, then 25 mM DTT was added. Digestion was initiated by addition of trypsin at a 50:1 (protein:enzyme) mass ratio. Samples were incubated at R.T. overnight. The reaction was then quenched by addition of trifluoroacetic acid (TFA) (final 2.5%). Peptides were desalted using a SPE column. The cartridge was primed (300 µL ACN), equilibrated (300 µL of 0.1% TFA in water), loaded twice, and washed (300 µL 5% ACN, 0.1 % TFA in water). Peptides were eluted (300 µL of 50% ACN, 0.1% TFA in water) into a new tube and dried down using a speedvac and stored at -20°C until LC-MS/MS analysis.

#### Mass spectrometry and data analysis

The samples were analyzed using a nanoflow-HPLC (Thermo Scientific EASY-nLC 1200 System) coupled to an Orbitrap Fusion Lumos Tribrid Mass Spectrometer (Thermo Fisher Scientific inc.) in data independent acquisition mode. Digested peptides were recovered in buffer A (3.9% ACN, 0.1% formic acid in water). Reverse phase separation of the peptides was done with an Aurora Ultimate™ analytical column (25 cm x 75 µm ID with 1.7 µm media, IonOpticks). Peptides were eluted with a solvent B gradient (0.1% FA in 80% ACN) for 120 min. The gradient was run at 400 nL/min with analytical column temperature set at 45°C. DIA analysis was done as reported by Mehta et al. 2022 with some modifications [25]. Full scan MS^1^ spectra (350 - 1400 m/z) were acquired with a resolution of 120,000 at 200 m/z with a normalized AGC Target of 200% and a maximum injection time of 20 ms. MS^2^ was acquired in the linear ion trap, ACG target value for fragment spectra was set to 2000%. Twenty-eight 38.5 m/z windows were used with an overlap of 1 m/z. Resolution was set to 30,000 using a dynamic maximum injection time and a minimum number of desired points across each peak set to 6.

DIA data analysis was performed in the software Spectronaut (v17) using direct DIA analysis workflow using default settings. The database for the searches was the Uniprot mouse proteome (2021, 55,336 sequences) with the hAPP and hCD33 sequence added for the Tg lines. Trypsin/P was selected as the digestion enzyme with a maximum of two missed tryptic cleavages and the search was performed with a maximum false discovery rate of 1% for peptides. Carbamidomethylation (C) was added as fixed modification and, deamidation at N/Q, and M oxidation were set as variable modifications. Before pairwise comparison of the populations under study, protein abundance variability was corrected by normalizing abundance using a global approach on the abundance average. Ratios for the comparisons, CD33M versus Control, CD33m versus Control, CD33M versus CD33m, were generated and a t-test was applied, the list of candidates was generated for significant proteins with *p*-value ≤ 0.05 and fold change ≥ 1.5 for each comparison. GO analysis for the selected candidates was performed directly in Spectronaut, over and underrepresented GO entries with *p*-value ≤ 0.05 were considered significant. Volcano plots between paired groups (CD33M versus Control, CD33m versus Control and CD33M versus CD33m) were generated on Prism (v9) using the list of proteins on each individual comparison. The triplot was generated using excel by initially graphing the log2(ratio) of each comparison on each side of a triangle. The intersection of lines between the specific fold change value and the vertex at the opposite end of the triangle corresponds to the proteomic correlation between the populations compared. The location of each circle on the graph indicates the association of that protein to the specific comparison. Identifications closer to the vertex, correspond to proteins with higher association to that genotype, while identifications closer to the center of the triangle indicate similar levels of that protein in all comparisons. From the lists of candidates generated, proteins with increased abundance to the CD33m and CD33M populations were extracted and highlighted in the triplot (blue and red, respectively).

### Isolation of adult mouse microglia

Isolation of microglia was performed as described previously [26]. Briefly, mice were perfused with 15 mL of ice-cold HBSS buffer containing Actinomycin D (5 μg/mL) and Triptolide (10 μM). Brain was then extracted and stored in 10 mL storage buffer containing Actinomycin D (5 μg/mL), Anisomycin (27.1 μg/mL) and Triptolide (10 μM) at 4 °C. Minced brains were homogenized in ice cold storage buffer with a 5 ml syringe plunger through a 40 μm filter (Corning) under sterile conditions. The cell suspension was then transferred to a 15 mL tube and centrifuged for 5 min at 4 °C (300 rcf). After centrifugation, the pellet was collected and resuspended in 10 mL of ice cold 40% Percoll (Sigma) diluted in 1x (final) HBSS and centrifuged (30 min, 500 rcf, 4 °C). After carefully removing the Percoll layer containing myelin debris, microglia were pelleted. After centrifuging (5 min, 300 rcf, 4 °C), the collected cell pellet was washed with 10 mL of ice-cold 1x HBSS buffer. Samples were resuspended in 50 μl of ice-cold flow buffer (0.5% BSA, 1 mM EDTA, in 1x PBS, Sterile Filtered) containing antibodies targeting Cd11b (BV510, clone ICRF44, Biolegend), CD45 (APC/Cy7, clone 30-F11, Biolegend), and Cx3cr1 (PerCP/Cy5.5, clone SA011F11, Biolegend) from Biolegend all at 1:200 dilution for 20 min on ice. Following incubation, the samples were washed with ice cold flow buffer, centrifuged (5 min, 300 rcf, 4 °C), and resuspended in 700 μl of ice-cold cell sorting buffer (1x HBSS containing 10% FBS and 1 mM EDTA) in preparation for cell sorting. An estimated 70,000 microglia (CD11b^+^, CD45^+^, Cx3cr1^+^) were sorted using the 100 μm nozzle at a sorting speed of approximately 3500 events/sec. We collected sorted samples in 1.7 ml Eppendorf tubes. Sorted cells were centrifuged (300 rcf, 5 min, 4 °C) and the supernatant was removed. Pelleted cells were resuspended in 100 μL PBS + 0.1% BSA and 10μL of the sample immediately counted using a Neubauer chamber using 0.4% Trypan blue solution (Thermo Fisher). For experiments, samples with a viability of over 95% were used. Cells were resuspended in PBS in an appropriate volume to achieve a concentration of 1000 cells/mL. This cell suspension was used to generate the gel-beads + cell emulsion by the 10X Chromium Controller (PN-1000202) using the Chromium Next GEM Single Cell 3′ GEM, Library & Gel Bead Kit v3.1 (PN-1000121), Chromium Next GEM Chip G Single Cell Kit (PN-1000120) and Single Index Kit T Set A, (PN-1000213). Reverse transcription, cDNA amplification, library preparation, and sample barcoding were performed following the available manufacturer’s protocol. Finally, sample libraries were pooled and sequenced in Illumina HiSeq P150 (Sequencing type: Paired-end, single indexing) to an average depth of ∼ 50,000 reads per cell.

#### Single-cell RNA sequencing

##### Library preparation

The FACS-isolated cells were processed on the 10x chromium controller following the 10x Genomics Next GEM Single Cell 3’ GEM, library, and v3.1 Gel Bead kit (10x Genomics; Cat. No. 1000121) and sequenced the samples on the Illumina HiSeq P150 sequencer at Novogene corporation inc. The samples were paired-end, single index sequenced at an average read depth of 50,000 reads per cell. The resulting BCL files were demultiplexed into FASTQ files and aligned to a custom *Mus Musculus* 10 (MM10) reference genome, adjusted to include *Clec7a*, a polymorphic pseudogene. The *Clec7a* annotation was added to the 10x Genomics pre-built MM10 genome (2020-A) GTF file from the 10x Genomics parent GTF file (gencode.vM23.primary_assembly.annotation.gtf.gz). The final custom genome was created by combining the MM10 genome (2020-A) FASTA file with the *Clec7a* modified GTF file using the cellranger mkref function in the 10x cell ranger pipeline (v3.0.0). Finally, the samples were aligned the custom genome using the cellranger count function to generate barcoded and sparse matrices, both raw and filtered, along with BAM files for downstream analyses.

##### Quality control, dimensionality reduction and clustering

Quality control, dimensionality reduction and initial clustering was performed in the R statistical environment (v4.1.2) using Seurat (v4.0; https://github.com/satijalab/seurat). A Seurat object was created per dataset to include genes expressed in a minimum of 3 cells and cells expressing a minimum of 200 genes using the CreateSeuratObject() function. The object was further refined to remove doublets and multiplets by removing cells with high gene counts (>3000 genes) and dead cell by removing cells high percentages of mitochondrial genes (>10%). All datasets were merged using the merge() function and normalized using the SCTransform() function according to the binomial regression model (Highly variable features = 3000, nCount and mitochondrial genes regressed).

Dimensionality reduction was performed using RunPCA(), FindNeighbors() (Dimensions = 15) and FindClusters() functions. 25 PCs were used for downstream analyses as determined by the PCA elbow plot. The FindClusters() function was run at multiple resolutions, ranging from 0 to 1, separated by 0.1. All clustering resolutions (0 to 1) were plotted on a tree generated by the Clustree package using the clustree() function. The 0.5 resolution was chosen for clustering based on the most stable level as identified by clustree().

The final clustering of the dataset was performed in a Jupyter notebook (v6.0.3) running a python environment (Python 3.8.3) and using Single Cell Clustering Assessment Framework (SCCAF) package (v0.0.10, https://github.com/SCCAF/sccaf). The SeuratObject was converted to an h5ad file using SeuratDisk by first converted the RDS file to an h5Seurat file using the Saveh5Seurat() and converting the h5Seurat file to an h5ad file using the convert() function (Destination = h5ad). The h5ad file was read into the python environment using Scanpy (v1.6.0)[27]. The clustering was refined using the SCCAF_optimize_all() function (minimum accuracy = 95%, iterations = 150). The machine learning algorithm iteratively clustered the dataset until a 95% self-projection accuracy was reached. The final clustering iteration was projected onto a UMAP using the sc.pl.umap() function.

### Open field assay

All mice were habituated in the behavior suite for 1 hr prior to testing. White noise was played via a sound device during habituation and during the experiment. Two white cube-shaped (40 cm x 40 cm) arenas, placed side-by-side were used for this experiment, enabling recording of two mice at a time. One mouse was placed in each arena for testing. Each mouse’s behavior and movement in a 15 min period were recorded and tracked via a camera connected to EthoVision 17 (Noldus, Wageningen, the Netherlands). Tracking was dependent on the mouse’s center-point as detected by EthoVision 17, with the nose- and tail-points being defined too. The center zone was defined in EthoVision 17 as a 30 cm x 30 cm area in the middle of the arena. After testing, the mouse was placed back into its home cage, and the arenas were thoroughly cleaned with 70% ethanol prior to testing the next mouse. Differences between the groups were tested with one-way ANOVA, followed by Tukey’s Test.

### Statistical analyses

Data represented as mean ± SD. The D’Agostino-Pearson normality test was used to test for Gaussian distribution of datasets. Differences between the groups were evaluated with one-way Anova followed by the Holm–Sidak Test. A probability of *P*<0.05 was considered indicative of significant differences between groups.

## Results

### Human CD33 isoforms distinctively alter Aβ plaque burden

To study the impact of hCD33 isoforms on the accumulation of Aβ *in vivo*, our previously developed STOP^flx/flx^-CD33M [19] and STOP^flx/flx^-CD33m [15] Tg mice were crossed with 5XFAD mice [20]. We previously established that these models give physiologically-relevant expression of hCD33 isoforms specifically in microglia when crossed with CX3CR1^Cre^ mice [15]. All experimental mice in this study had a single copy of CX3CR1^Cre^ to drive specific expression of the transgene in the microglia cell lineage and were aged to four or eight months. Because of the known sex-dependent differences in accumulated Aβ levels in the 5XFAD model, and to investigate sex-specific impacts from hCD33 isoforms on Aβ deposition, cohorts of both male and female mice were examined [20, 28, 29]. We first conducted immunohistochemistry analysis by fluorescence microscopy of coronal hemi-brain slices stained with an anti-Aβ antibody and quantified the area of deposited Aβ as a percentage of the total brain area (**Fig. 1**). At four months, there was no significant difference between the control and CD33M group, but CD33m mice had significantly reduced levels of Aβ compared to both control and CD33M mice (**Fig. 1a,b**). Trends were similar in both female and male mice and we could not detect any sex-specific differences between hCD33 groups compared to the control. However, female mice from all genotypes showed augmented deposited Aβ levels (**Suppl. Fig. 1a,b**), which is consistent with previous reports in 5XFAD mice [20, 30]. Also, in keeping with previous observations in the 5XFAD model, the dorsal subiculum showed a high Aβ plaque density [20, 31, 32]. At eight months, all cohorts presented significant differences in deposited Aβ levels, with CD33M mice having higher levels of Aβ compared to the other groups, while CD33m mice showed lowest Aβ levels (**Fig 1c,d**). Increased and decreased Aβ deposition in CD33M and CD33m mice, respectively, was evident in both sexes at eight months (**Suppl. Fig. 1c,d**). Due to the absence of any notable sex driven effects from either of hCD33 isoforms, we pursued the rest of the analyses in combined sexes.

**Figure 1:**
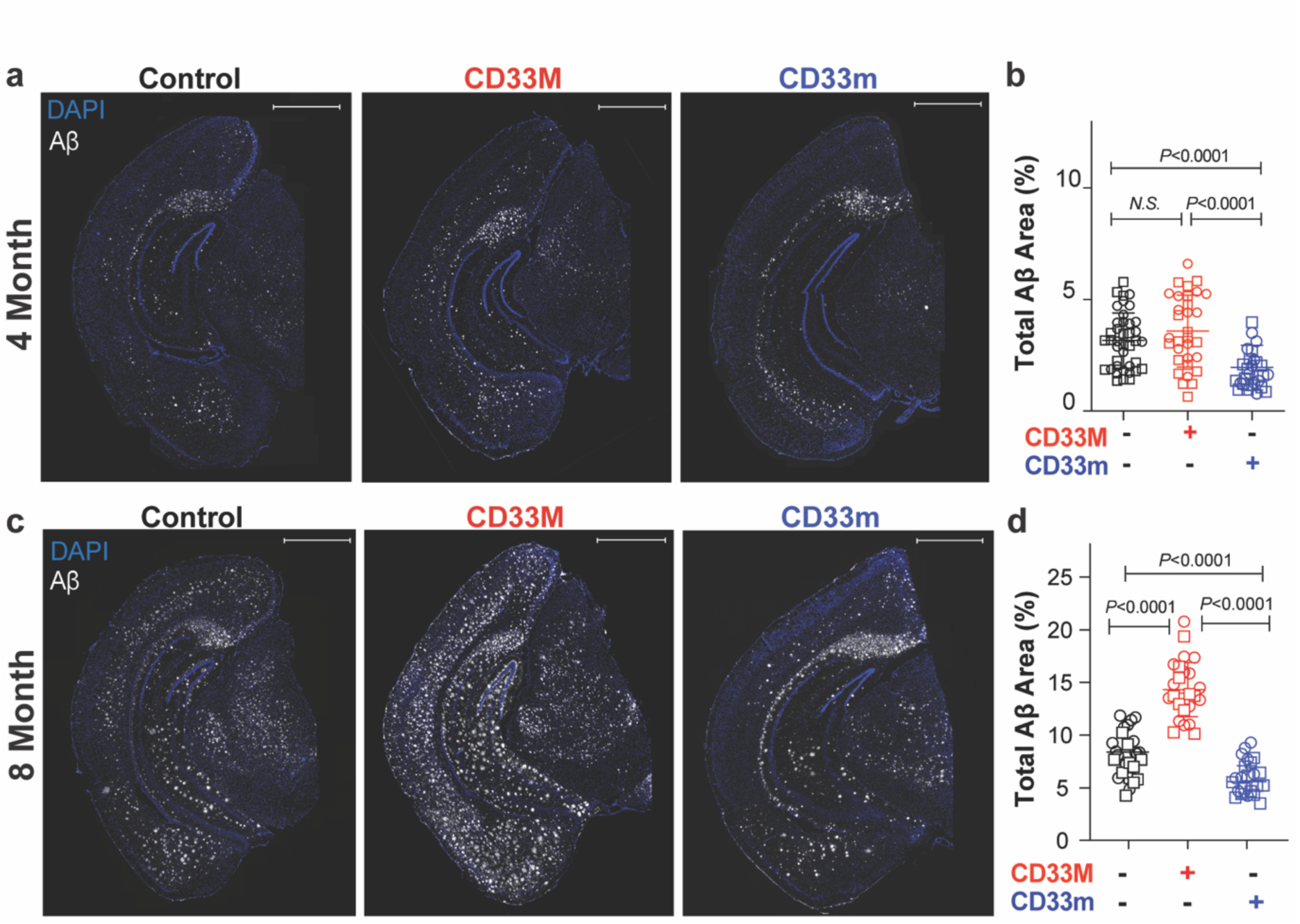
hCD33 isoforms have opposing effects on Aβ accumulation in 5XFAD mice. **a)** Representative epifluorescent images of Aβ deposition in 5XFAD mice at 4 months by IF imaging with the anti-Aβ antibody (clone MOAB2, white) and DAPI (blue). **b)** Quantification of total Aβ levels at 4 months in pooled male (squares; n = 20, 18, and 12 for control, CD33M, and CD33m, respectively) and female (circles; n = 20, 15 and 15 for control, CD33M, and CD33m, respectively) mice. **c)** Representative epifluorescent images of Aβ deposition in 5XFAD mice at 8 months. **d)** Quantification of total Aβ levels at 8 months in pooled male (squares; n = 19, 11, and 10 for control, CD33M, and CD33m, respectively) and female (circles; n = 22, 14 and, 13 for control, CD33M, and CD33m, respectively) mice. Scale bar = 1000 µm.

We also analyzed two additional parameters: region-specific analysis of deposited Aβ levels in cortex and hippocampus, and the number of Aβ clusters in whole brain. At four months of age, CD33m mice showed less Aβ deposition in the cortex compared to control and CD33M mice, but no differences between the groups were observed in the hippocampus at this timepoint (**Suppl. Fig. 1e,f**). At eight months, Aβ deposition was significantly different between all three groups in the cortex, while in the hippocampus the only significant difference was between CD33M versus control and CD33m mice (**Suppl. Fig. 1g,h**). Quantitative analysis of the number of Aβ clusters pointed to no significant differences between any of the groups at four months (**Suppl. Fig. 1i**). However, by eight months these differences reached statistical significance with CD33m mice showing overall fewer Aβ clusters and CD33M mice showing more Aβ clusters compared to the control 5XFAD mice (**Suppl. Fig. 1j**).

### Expression of hCD33 isoforms significantly alter Aβ plaque composition

Aβ deposits exhibit varying compositions, including dense core and diffuse morphologies (**Fig. 2a**). Dense core plaques are predominately composed of fibrillar Aβ aggregates and are readily detected with amyloid-binding dyes such as Thioflavin S (ThioS) [33–35]. In contrast, diffuse plaques, typically containing dispersed protofibrils and oligomers, stain weakly with ThioS but can be readily detected with Aβ antibodies. To investigate the impact of hCD33 isoforms on plaque composition, we stained brain slices with ThioS in parallel with an anti-Aβ antibody. Within the whole brain slice, CD33M mice had significantly fewer number of Aβ clusters with a ThioS core at both timepoints (**Fig. 2b-e**). Unexpectantly, CD33m had a significantly higher percentage of ThioS^+^ clusters compared to the other two groups at eight months **Fig. 2b-e**). These findings suggest that hCD33 isoforms impact more than just Aβ deposition and have strong effects on plaque composition. Specifically, these observations point to a shift in plaque composition towards a more compact form in the CD33m mice, which is the isoform associated with protective effects in the context of AD [11, 18].

**Figure 2:**
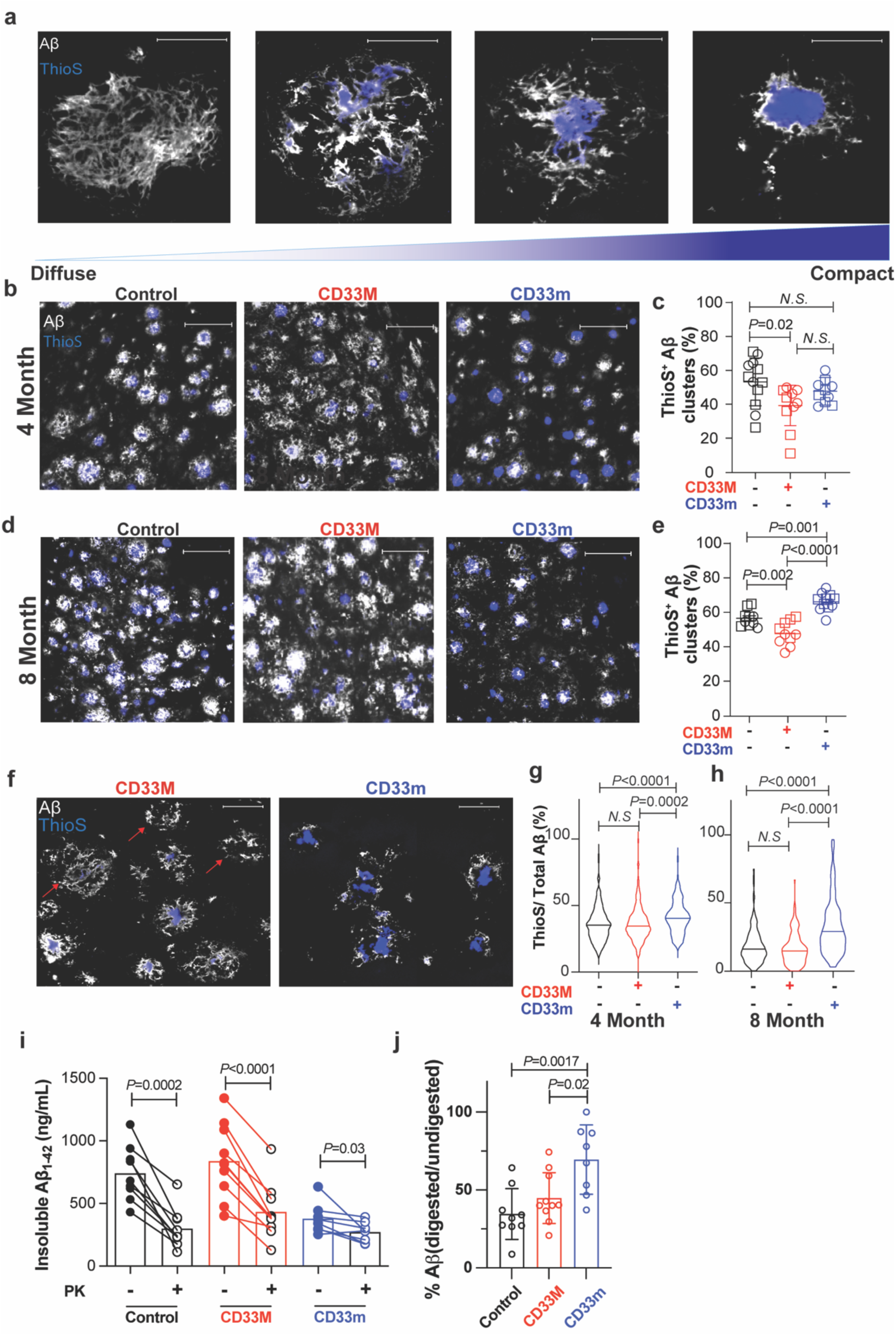
Altered Aβ plaque composition by hCD33 isoforms. **a)** Representative confocal fluorescent images showing the different degrees of plaque compaction by co-staining for total Aβ and ThioS. A diffused plaque without ThioS staining at the far left, a plaque with small ThioS core in the middle, and a highly compact plaque at the far right are presented. Scale bar = 20 µm **(b-e)** Quantification of Aβ clusters containing a ThioS^+^ core. Representative epifluorescent images of Aβ deposits in the dorsal subiculum of 5XFAD mice at **(b)** 4 and **(d)** 8 months, with co-staining with anti-Aβ antibody (white) and ThioS (blue). Scale bar = 50 µm. Quantification for pooled males (squares; n = 5 per genotype) and female (circles; n = 5 per genotype) mice at **(c)** 4 and **(e)** 8 months. **f)** Representative confocal fluorescent images of plaque composition in the dorsal subiculum of 8 months old CD33M and CD33m mice. Scale bar = 20 µm **(g,h)** Quantification of plaque composition (ratio of Aβ over ThioS levels) in the dorsal subiculum of 5XFAD mice at **(g)** 4 and **(h)** 8 months. A total of 300 plaques per mouse, from 5 males and 5 females, were quantified for each group. **i)** Biochemical characterization of PK-sensitivity of insoluble Aβ1-42 from 5XFAD mice. **j)** Quantification of the percentage of insoluble Aβ1-42 levels after PK digestion compared to not treated with PK.

To examine the Aβ plaque composition in greater detail, we carried out quantitative analyses of the dorsal subiculum and cortex at the eight-month timepoint. Consistent with the global analysis of the hemi-brain, we observed the highest deposited Aβ levels in CD33M mice compared to both the control and CD33m mice (**Suppl. Fig. 2a**). ThioS levels in CD33M mice were highest compared to the other two groups, however, CD33m mice also showed higher ThioS levels in the dorsal subiculum compared to the control group (**Suppl. Fig. 2b**). When examining the ratio of ThioS to the amount of total deposited Aβ, the results once again point to CD33m promoting more compact Aβ while CD33M promotes the opposite outcome (**Suppl. Fig. 2c**).

These differences in plaque composition motivated further analysis of individual plaques in dorsal subiculum and cortex (**Fig. 2f**). At four months, CD33m mice had smaller Aβ clusters than other groups, while CD33M and control showed no differences (**Suppl. Fig. 2d**). However, CD33M had larger ThioS cores (**Suppl. Fig. 2e**). As a result of smaller Aβ clusters, despite no decrease in the size of ThioS core, CD33m promoted higher plaque compaction compared to CD33M and control (**Fig. 2g**). At eight months, both CD33M and CD33m had a significant impact on the size of Aβ clusters, with CD33M promoting larger Aβ cluster size and CD33m promoting smaller Aβ cluster size (**Suppl. Fig. 2f**). Despite an overall smaller Aβ clusters, CD33m promoted larger ThioS core area in the plaques compared to the control (**Suppl. Fig. 2g**). Therefore, CD33m promoted the highest level of plaque compaction compared to other genotypes, while CD33M showed no significant difference compared to the control group (**Fig. 2h**). It is worth noting that for the analyses on individual clusters, we focused on ThioS^+^ plaques, whereas the global plaque compaction in the dorsal subiculum region did not differentiate between the plaques with or without a ThioS core.

To perform further biochemical analysis on Aβ deposits, we extracted soluble and insoluble Aβ from the snap-frozen half brain of 5XFAD mice at eight months and quantified Aβ1-42 levels by electrochemiluminescence. While soluble Aβ1-42 levels were not significantly different between the genotypes (**Suppl. Fig. 3a**), insoluble levels were significantly lower in CD33m mice compared to the other two groups (**Suppl. Fig. 3b**). As different forms of Aβ aggregates show different sensitivity to protease digestion [36, 37], we assessed the sensitivity of the insoluble fraction to proteinase K (PK) digestion [38]. A titration of PK was first performed to optimize the concentration that produced the maximum decrease in Aβ levels in the insoluble fraction (**Suppl. Fig. 3c**). Accordingly, a PK concentration of 20 µg/ml was selected and applied to samples from all genotypes, which led to a significant decrease in Aβ1-42 levels in all groups (**Fig. 2i**). The remaining levels of Aβ1-42 post-digestion was indistinguishable between all groups, which was likely due to the presence of highly compacted, PK-resistant material. However, the percentage of Aβ1-42 relative to the undigested control in each group demonstrated that CD33m mice had significantly less PK-sensitive Aβ1-42 compared to the others (**Fig. 2j**). Taken together, the combination of these analyses further supports our earlier observations, suggesting that CD33m expression in microglia leads to decreased total Aβ levels and more specifically lower levels of diffuse, PK-sensitive aggerated Aβ assemblies.

### hCD33 isoforms modulate microglial cell response to Aβ deposits

Microglial cell phagocytosis *in vitro* is increased by CD33m and decreased by CD33M [15, 16, 19, 39]. As a proxy for phagocytosis *in vivo*, we quantified the amount of internalized Aβ in microglia by co-staining tissue sections with anti-Aβ and anti-IBA-1 (**Fig. 3a**) [40]. At eight months, decreased levels of internalized Aβ were observed in CD33M microglia, while CD33m microglia had increased levels of internalized Aβ compared to control 5XFAD microglia (**Fig. 3b**). The decreased and increased levels of internalized Aβ in CD33M and CD33m microglia respectively, could explain the overall trends observed in Aβ levels, plaque compaction, and brain plaque load. We next analyzed the peri-plaque microglial cells, also known as plaque-associated microglia (PAM), in greater detail by confocal microscopy (**Fig. 3c**). Compared to control 5XFAD mice, CD33M mice had decreased numbers of PAM, while CD33m had increased numbers of PAM at eight months (**Fig. 3d**). These differences were also observed at four months (**Fig. 3e**). These trends were not due to a global effect on the number of microglia cells, as overall IBA-1 levels at eight and four months showed the opposite trend and revealed increased IBA-1 density in CD33M mice compared to other groups (**Suppl. Fig 4**). Of note, the increased IBA-1 density could reflect on microgliosis in the CD33M mice or increased inflammatory response due to greater plaque burden.

**Figure 3:**
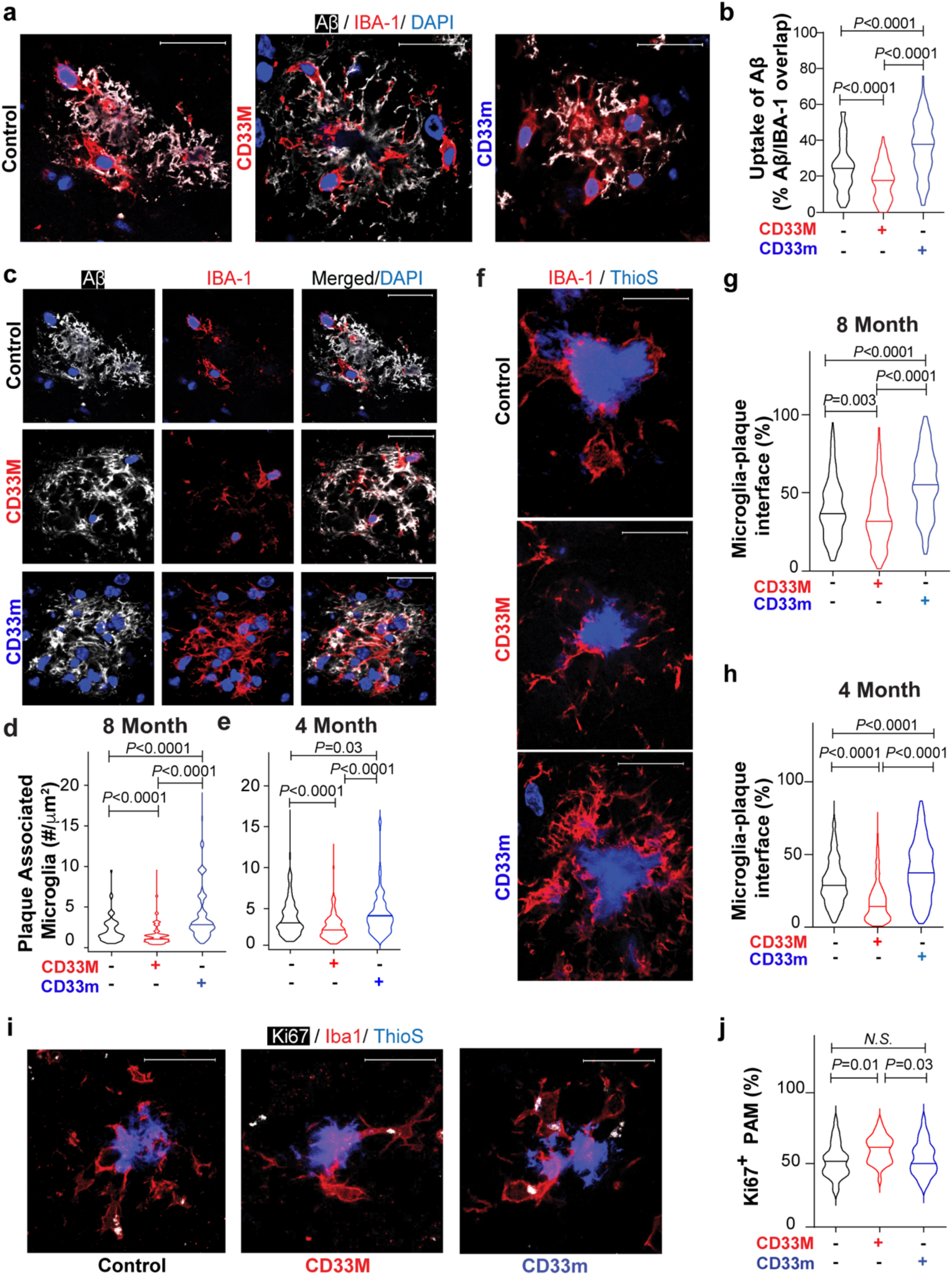
hCD33 isoforms modulate differential response of microglia to Aβ. **a)** Representative confocal fluorescent images of internalized Aβ by microglia of control, CD33M, or CD33m 5XFAD mice at 8 months. IF images are co-stained with anti-Aβ antibody (white), anti-IBA-1 (red), and DAPI (blue). Scale bar = 20 µm. **b)** Quantification of internalized Aβ by microglia measured as the area of Aβ signal overlapping with IBA-1. A total of 300 Aβ clusters per mouse, from 5 males and 5 females, were quantified for each group. **c)** Representative confocal fluorescent images of plaque associated microglia in control, CD33M, and CD33m mice. IF images are co-stained with anti-Aβ antibody (white), anti-IBA-1 (red), and DAPI (blue). Scale bar = 20 µm. **(d,e)** Quantification of plaque associated microglia normalized to µm^2^ of plaque area. A total of 300 plaques, in 5 males and 5 females, per group were analyzed at **(d)** 8 and **(e)** 4 months. **f)** Representative confocal fluorescent images of microglia-ThioS core interface in control, CD33M, and CD33m mice. IF images are co-stained with anti-IBA-1 (red), and ThioS (blue). Scale bar = 20 µm. **(g,h)** Quantification of plaque-microglia interface in 5XFAD mice at **(g)** 8 and **(h)** 4 months, measured as the percentage area of ThioS perimeter with overlapping IBA-1 signal. A total of 300 plaques, in 5 males and 5 females per genotype, were analyzed. **i)** Representative confocal fluorescent images of Ki67 staining of PAM in 5XFAD mice at 8 months in CD33m and CD33m mice. IF images are co-stained with anti-ki67-1 (white), anti-IBA-1 (red), and DAPI (blue). Scale bar = 20 µm. **j)** Quantification of Ki67^+^ PAM measured as the percentage of PAM with KI67 punctate inside the cell. PAM of 300 plaques, from 5 males and 5 females per genotype, were analyzed.

As microglia promote compaction through biochemical and biophysical mechanisms [40–42], we analyzed the microglia-ThioS core contact sites by quantifying the perimeter of the ThioS core interface that was covered by IBA-1 signal (**Fig. 3f**). We observed a significant increase in microglia-plaque interface in CD33m mice at both eight months (**Fig. 3g**) and four months (**Fig. 3h**). Accordingly, CD33M mice showed the opposite trend regarding microglia-plaque interface. These results reveal another way that hCD33 isoforms modulate microglia response to Aβ deposits. Finally, increased numbers of PAM in the CD33m group, despite their lower global IBA- 1 density, prompted us to investigate the origins of these effects. We started by quantifying the proliferative capacity of PAM at eight months old mice through assessing the proliferative marker Ki67^+^ within PAM (**Fig. 3i**). In line with previous observations, approximately 50% of PAM were Ki67^+^ [43, 44], however, CD33m had no effect on the percentage of Ki67^+^ PAM compared to the control. Interestingly, CD33M-expressing microglia had a modest yet significant increase in percentage of Ki67^+^ PAM compared to other groups, which may be in line with the higher IBA-1 density in these mice (**Fig. 3j**). Regardless, these results rule out enhanced proliferation as a mechanism for enhanced numbers of PAM in CD33m mice, suggesting that other mechanisms may be at play.

### A protein implicated in cell migration is upregulated in CD33m^+^ microglia

We next took an unbiased approach to identify proteins that may contribute to the enhanced response of microglia to Aβ plaques in CD33m mice. Accordingly, we applied label-free mass spectrometry to perform global quantitative proteomics on brain homogenates of mice at eight months (**Fig. 4a**). More than 6000 proteins were consistently identified in each biological replicate by our data-independent acquisition mass spectrometry approach (DIA-MS). On average, 6697 proteins were found in the CD33M mice, 6727 for the CD33m mice, and 6732 for the control. The abundance of all those proteins was quantified in all samples and reported here (**Suppl. File 1**). Examining differentially expressed proteins revealed numerous differences in global proteome of CD33M or CD33m-expressing microglia compared to the control group (**Suppl. Fig. 5**). Significantly increased and decreased proteins were defined as those with a fold change larger than 1.5 or smaller than 0.66 respectively, and a *p-*value ≤ 0.05. Comparison of CD33M versus control pointed to a total of 47 proteins significantly increased in the CD33M brain, while 96 proteins were decreased. Gene ontology (GO) enrichment analysis of the significantly changed proteins (**Suppl. Fig. 6**) revealed alterations in pathways involved in inflammatory response, defective lipid metabolism, increased complement-dependent cytotoxicity, and cellular oxidative stress. The second comparison was done on CD33m versus control, wherein 40 proteins showed a significant increase in the CD33m, and 235 proteins were decreased compared to the control. Among the significant GO entries for the protein candidates, increased phagocytosis/engulfment, enhanced neuronal health and innervation, and improved lipid synthesis and storage were among the top biological processes with alterations. Finally, for the third comparison, CD33M versus CD33m, 17 increased proteins and 122 decreased proteins showed significant changes. Decreased phagocytosis or engulfment, along with disturbed cell adhesion, nerve morphogenesis and neuronal remodeling were consistently identified within the altered biological processes.

**Figure 4:**
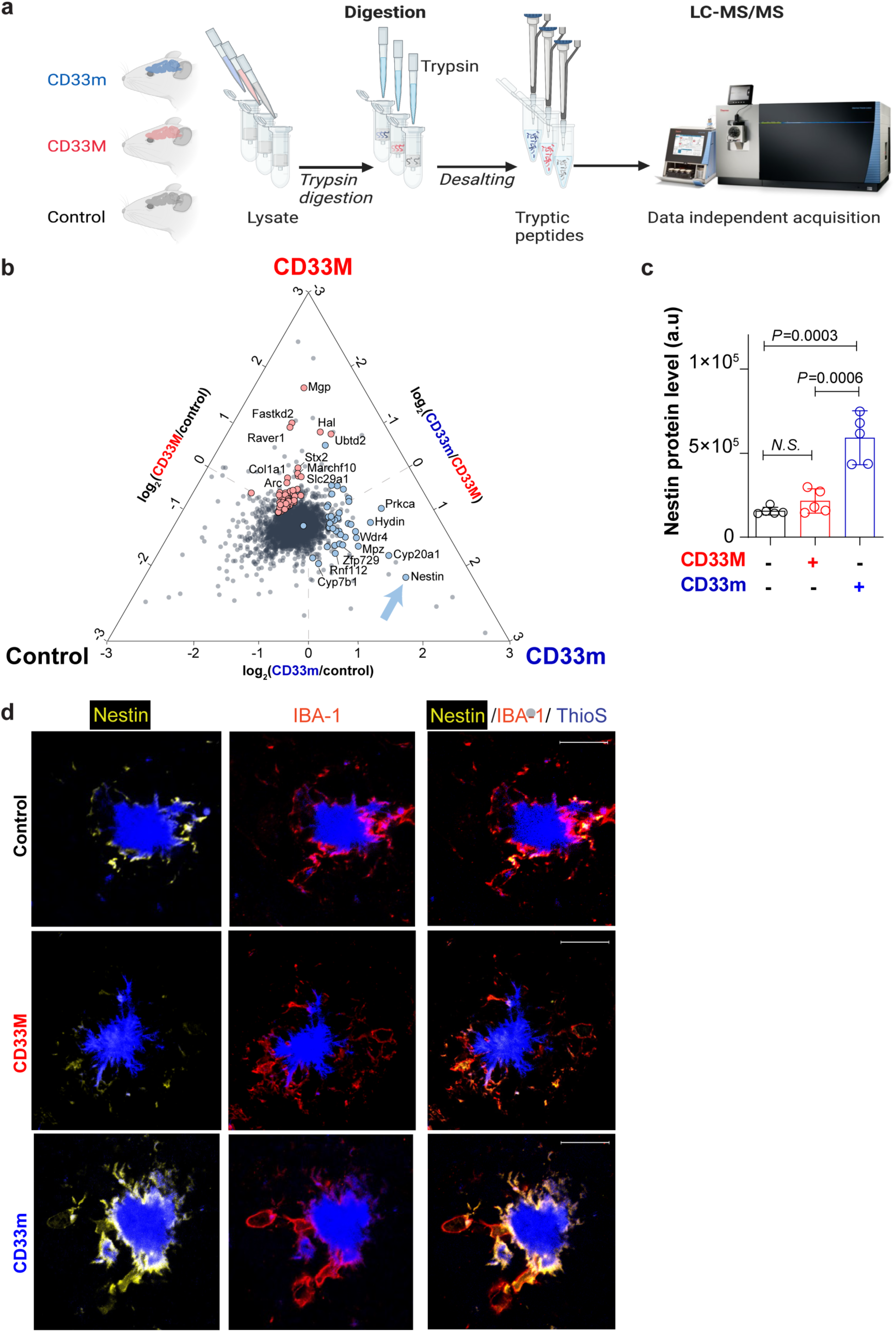
Quantitative proteomics reveal changes in protein abundance in the brain of 5XFAD mice expressing hCD33 isoforms. **a)** Schematic representation of the mass spectrometry workflow used for the assessment of global proteome changes in the brain of 5XFAD mice expressing CD33M or CD33m compared to control at 8 months. A total of 5 male mice were used per genotype. On-column digestion and data-independent acquisition mass spectrometry (DIA-MS) was used to quantify protein levels. **b)** Correlation of the proteomic changes in mouse brain for all proteins identified by DIA-MS. The triplot shows protein abundance changes calculated from each pairwise comparisons (CD33M vs control, CD33m vs control, and CD33m vs CD33M). Identifications closer to the vertex correspond to proteins with higher association to that genotype, while identifications closer to the center of the triangle indicate similar levels of that protein in all comparisons. Proteins with significant changes (fold change ≥ 1.5 and *p*-value ≤ 0.05) are highlighted. The proteins increased in mouse brains expressing CD33m are colored in blue, while the proteins increased in the CD33M isoform are colored in red. **c)** Quantification of nestin in each individual sample as measured by mass spectrometry. **d)** Representative confocal fluorescent images of nestin staining of PAM in 5XFAD mice at 8 months in control, CD33m, and CD33m mice. IF images are co-stained with anti-nestin (yellow), anti-IBA-1 (red), and ThioS (blue). Scale bar = 20 µm.

Correspondence of the proteomic changes among populations was evaluated by correlating the fold change between comparisons using a triplot. The graph generated shows the protein abundance distribution measured using DIA-MS and its association with the mice genotypes (**Fig. 4b**). Within the proteins with high association to CD33m mice, nestin was identified as the top hit with significant changes in both comparisons CD33m versus control and CD33M versus CD33m. Assessing the proteins with different levels that were associated with enhanced phagocytosis in CD33m mice revealed increased levels of nestin in the comparisons CD33m versus control and CD33M versus CD33m (**Fig. 4b,c**). Nestin is an intermediate filament associated with cell mobility/migration and has been reported to be upregulated in microglia under specific circumstances including activation states [45]. Further immunostaining analysis of brain slices with an anti-nestin antibody clearly demonstrates that nestin protein levels were uniquely elevated in microglia-contact sites in CD33m mice at eight months (**Fig. 4d**). These results further support our earlier observations, regarding a higher density of PAM and enhanced microglia-plaque interface in CD33m.

### Transcriptional profiling of microglia supports distinct effects of hCD33 isoforms

To further explore the principles contributing to differences between hCD33 isoforms in modulating microglia response, we carried out single-cell RNA sequencing (scRNAseq) analysis on microglia of 5XFAD mice from all three genotypes at eight months, along with one age-matched control non-5XFAD. After the quality control measures were carried out, the four datasets were pooled and clustered with Seurat V5, and single cell clustering assessment framework (SCCAF) was applied to re-cluster cells based on classic microglial gene expression of *Hexb, Fcrls, Sall1, Tmem119*, and *P2ry12*. Overall, we obtained 21,714 cells, which was reduced to 15,200 after quality control. Clustered cells were projected onto a uniform manifold approximation and projection (UMAP) to identify fourteen clusters, falling into six main categories: border associated macrophages (BAM), homeostatic microglia (HM), transitioning microglia (TM), RNA binding protein microglia (RBM), myelin transcript enriched microglia (MTEM), and disease associated microglia (DAM) (**Fig. 5a**). The four groups of mice (**Fig. 5b-e**) had a different distribution of cells in each of the subsets (**Fig. 5f,g**). Proliferative microglia expressed classic proliferative genes such as *Mki67* and *Top2a* (**Fig. 5h**). BAM are found in the meninges and perivascular space express markers such as *Mrc1* and *Ms4a7* [46–48]. We identified three, highly related, HM clusters enriched in 5XFAD CD33M. Adjacent to HM we found two clusters of microglia we referred to as TM, as they expressed higher levels of Trem2 than HM microglia. MTEM expressed non-microglial, oligodendrocyte lineage cell transcripts *Mbp* and *Plp1,* consistent with oligodendrocyte renewal in this model [49].

**Figure 5:**
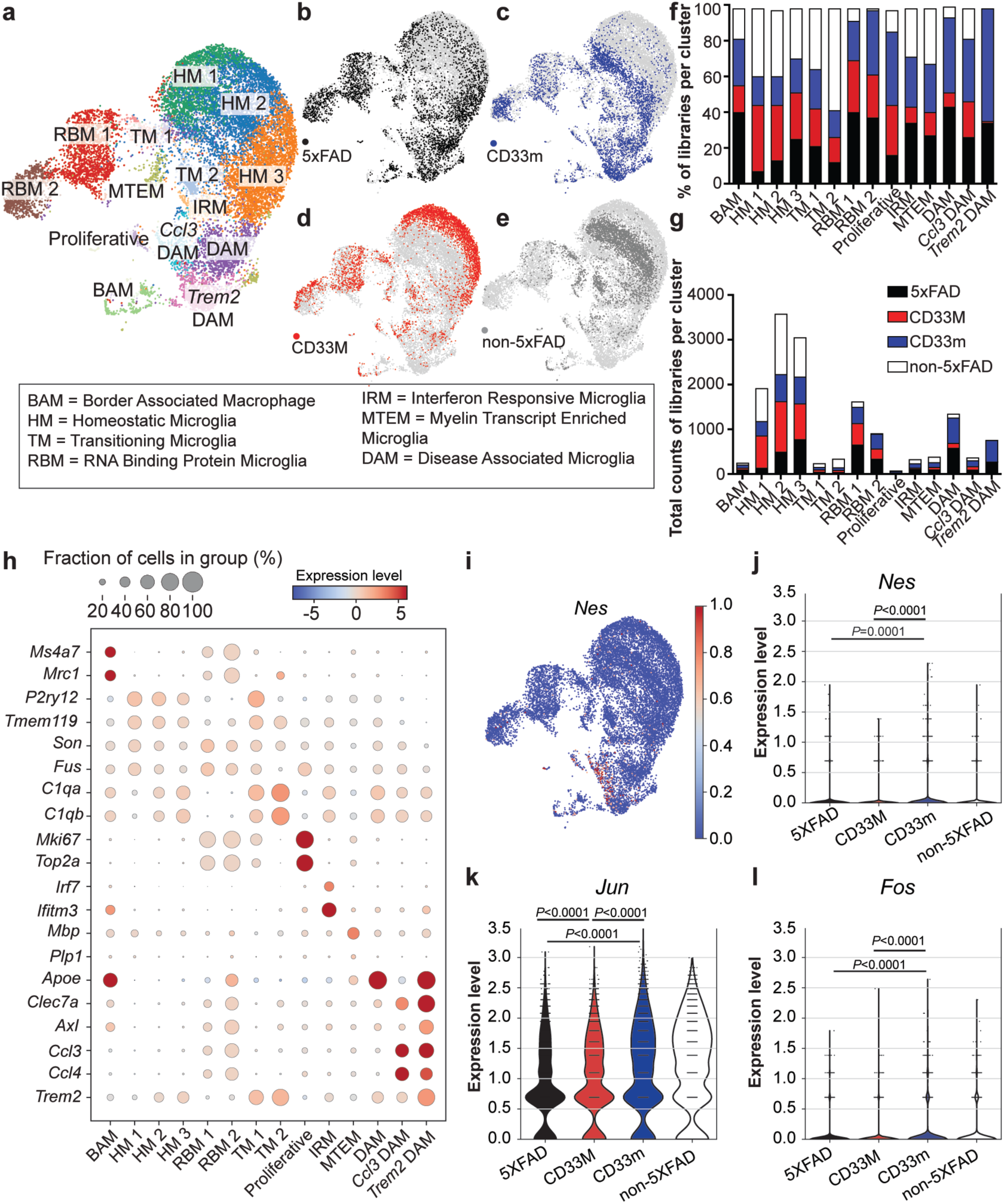
Single-cell RNA sequencing reveals *Ccl3* and *Trem2* DAM enriched in the CD33m genotype. **a)** Unsupervised and iterative machine-learning based clustering of 15,200 microglia (*Hexb*^+^*Fcrls*^+^*Tmem119*^+^*Sall1*^+^) and BAM (*Ms4a7*^+^*Mrc1*^+^*Lyve1*^+^*Timd4*^+^) collected from control 5XFAD, CD33M 5XFAD, CD33m 5XFAD, and control non-5xFAD mice. Microglia from three homeostatic (HM1-3), two transitioning (TM1-2), two RNA binding protein (RBM1-2) subpopulations along with the disease-enriched interferon responsive (IRM), myelin transcript enriched (MTEM), and disease associated (DAM) clusters. **(b-e)** UMAPs of individual samples showing **(b)** control 5XFAD, **(c)** CD33m 5XFAD, **(d)** CD33M 5XFAD, and **(e)** control non-5xFAD. **(f,g)** Separation of each cluster by **(f)** proportion of cells and **(g)** absolute number of cells belonging to each genotype. DAM, specifically *Trem2* DAM, are enriched in the CD33m genotype and reduced in the CD33M. **h)** Differential gene expression per cluster used to define microglial subpopulations. HM are characterized by homeostatic genes (*P2ry12, Tmem119*), RBM by genes related to RNA binding (*Son, Fus*), TM by a combination of homeostatic (*P2ry12, Tmem119*), complement (*C1qa, C1qb*), and proliferative (*Mki67, Top2a*) genes. DAM are defined by *Clec7a* and *Apoe* expression and further delineated by expression of *Ccl3* and *Trem2*. **i)** UMAP showing enriched *Nes* expression within the *Ccl3* DAM cluster. **(j-l)** Violin plots showing the upregulation of **(j)** *Nes*, **(k)** *Jun*, and **(l)** *Fos* in the CD33m 5XFAD group relative to control 5XFAD and CD33M 5XFAD.

We identified three different DAM subsets based on the expression of classic DAM markers *Clec7a* and *Axl,* which were absent in the non-5XFAD group (**Fig. 5f,g**). In addition to these classic DAM markers, we identified a DAM cluster specifically enriched for the chemokines *Ccl3 and Ccl4.* These factors were recently described to be released by microglia and act on neuronal Ccr5 to disrupt autophagy [50]. CD33m microglia were enriched within *Ccl3* DAM, and a DAM population enriched for these genes, but also expressing heightened levels of *Trem2* (termed *Trem2* DAM). The enrichment of CD33M microglia in HM clusters and reduction of CD33M microglia in DAM clusters is reminiscent of mice lacking TREM2 [51]. Given that DAM express genes related to phagocytosis such as *Cd86*, *ApoE*, *Lpl*, and *Cst7,* these data are consistent with heightened phagocytosis in CD33m^+^ microglia. To follow up these findings, we specifically examined the expression of *Clec7a*, which is upregulated in DAMs (**Suppl. Fig. 7a**) by IF staining of plaques and found that there was a significantly higher level of Clec7a in CD33m mice and a significantly lower level in CD33M mice (**Suppl. Fig. 7b,c**). Nestin gene (*Nes*) expression was specific to DAM (**Fig. 5i**) and, in line with the proteomics data, its transcript levels were statistically higher in CD33m microglia (**Fig. 5j**).

We also identified two clusters of microglia enriched for RNA binding proteins (termed RBM1 and RBM2), such as *Fus* and *Son*. RBM1 and RBM2 expressed low levels of DAM markers, suggesting they are DAM-like, while RBM2 expressed higher levels of *Apoe*, indicating this was a more reactive microglial population. We found no overt differences between CD33M and CD33m in RBM. Moreover, consistent with our previous report in homeostatic conditions [15], we observed upregulation of the immediate early genes (IEGs), such as *Jun*, and *Fos*, in 5XFAD-CD33m microglia (**Fig. 5k,l**). Fos is recruited to enhancers of scavenging and pathogen recognition receptors in microglia [52], which is consistent with *Fos* upregulation representing a more vigilant state.

### hCD33 isoforms distinctively affect neuronal health and behavior

The substantial impact of hCD33 isoforms on total Aβ levels, as well as plaque composition motivated us to further explore their effects on neuronal health and neurobehavior of the mice. GO enrichment analysis gave a hint of enhanced neuronal health in CD33m mice compared to control **(Suppl. Fig. 6)**. Additionally, we analyzed dystrophic neurites (DNs) by staining for LAMP1, a marker of lysosomal vesicles in swollen neuritic plaques [53, 54]. Quantification of DNs in the brain subiculum and cortex further supported the detrimental and protective effects of CD33M and CD33m, respectively. CD33M mice had significantly higher levels of dystrophic neurites while CD33m mice had lower levels compared to the control group (**Fig. 6a,b**). Quantification of individual DNs revealed that CD33M mice had on average larger DNs compared to the two other groups (**Fig. 6c,d)**. Moreover, the smaller neuritic plaques in CD33m had a higher number of PAM compared to both control and CD33M groups (**Suppl. Fig. 8a,b**).

**Figure 6:**
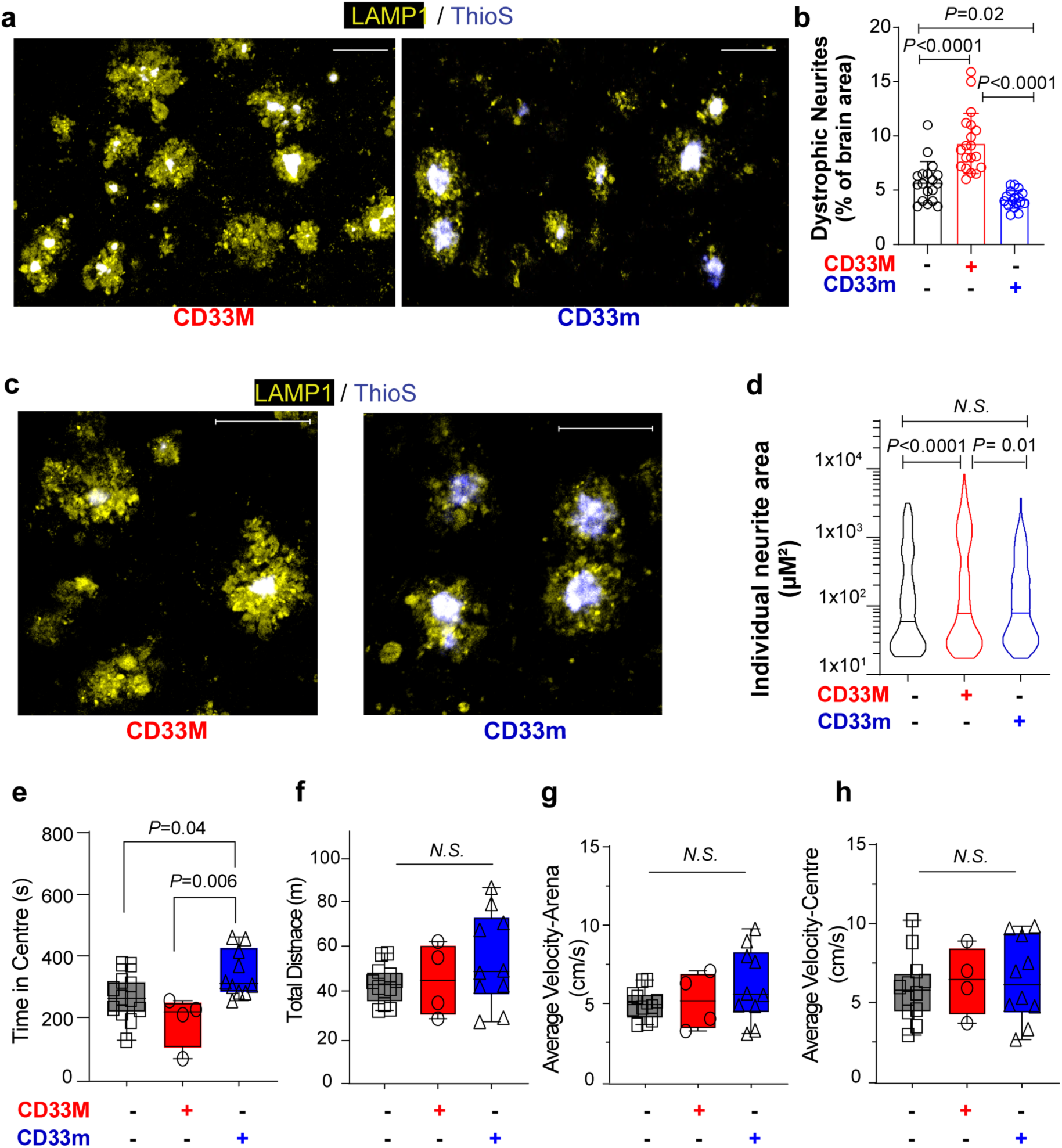
hCD33 isoforms impact neuronal health and animal behavior. **a)** Representative immunofluorescent confocal images of dystrophic neurites stained with lysosomal marker LAMP1 (yellow) in the dorsal subiculum of 5XFAD CD33M and CD33m mice at 8 months. Scale bar = 50 µm. **b)** Quantification of total area of DNs in the dorsal subiculum. A total of 5 male and 5 female mice per genotype were used. **c)** Higher magnification immunofluorescent confocal images of individual DNs in the dorsal subiculum of 5XFAD CD33M and CD33m mice at 8 months. Scale bar = 50 µm. **d)** Quantification of DN area in individual neuritic plaques of 5XFAD control, CD33M, and CD33m mice at 8 months. A total of 500 plaques from 5 males and 5 females per genotype were quantified. **(e-h)** Results from an open field assay carried out on 5XFAD mice at 1 year. Quantification of **(e)** the time mice spent in the central region of the field, **(f)** total distance travelled, **(g)** average velocity in the whole arena, and **(h)** average velocity in the central region.

Additionally, we performed an open field test on 5XFAD mice at 1-year of age [55]. This behavioral test has been previously used to identify altered cognitive performance associated with anxiety and exploratory behavior in 5XFAD mice [56]. As all our datasets pointed to sex-independent effects of hCD33 isoforms, we focused on female mice with more aggressive pathology and cognitive deficits in behavioral traits [57, 58]. CD33m mice spent significantly more time in the center compared to other groups (**Fig. 6e**), suggesting decreased anxiety and improved neurobehavioral outcomes in these animals compared to CD33M and control group. All other parameters, such as total distance and average velocity in the whole arena and the center (**Fig. 6f**-h), were similar between all groups.

## Discussion

Numerous studies have linked polymorphisms in *CD33* to AD [8, 9, 59, 60], yet the exact function of the expressed protein in microglia is poorly understood. Studying the role of hCD33 *in vivo* is confounded by the fact that mCD33 is functionally divergent compared to hCD33 and, more importantly, does not generate an equivalent CD33m protein isoform [18, 19, 61]. Accordingly, mouse models of AD miss the CD33 component of human-specific microglial cell biology. Herein, leveraging Tg mice expressing either of the hCD33 isoforms in the microglial cell lineage, we explored the impact of each isoform and discovered profound and opposite effects of hCD33 isoforms in the context of aggressive amyloid deposition. Specifically, CD33M, preferentially encoded by the rs12459419C AD-susceptible allele in humans, exacerbated Aβ deposition and resulted in decreased plaque compaction. Conversely, CD33m, preferentially encoded by the rs12459419T AD-susceptible allele, decreased Aβ deposition and increased plaque compaction. These results point to a combination of loss-of-function and gain-of-function effects delivered by CD33m. Accordingly, there are two beneficial effects that come from the rs12459419T *CD33* allele: decreased CD33M expression and increased CD33m expression [62]. Based on a CD33-null allele (rs201074739) lacking any AD-protective effects [14, 63], the loss-of-function alone may be insufficient to produce an AD-protective effect, suggesting the gain-of-function role is at play for beneficial effects of rs12459419T.

Differences in CD33m mice, manifested as decreased total Aβ levels, enhanced plaque compaction, and higher number of PAM were consistently significant at both four and eight months timepoints. In contrast, changes in Aβ deposition and microglial cell response in CD33M took longer to develop, reaching significance only at eight months. Accordingly, in our models the overall impact of CD33m on modulating microglia and Aβ deposition is stronger and more profound compared to CD33M. This is in agreement with our previous work, wherein we reported that the *in vitro* gain-of-function role of CD33m is dominant over the suppressive effect of CD33M [15]. Regardless, the timing of events in our datasets reinforce the idea that hCD33 isoforms play unique and divergent roles in microglia.

CD33M mice presented higher levels of deposited Aβ compared to other genotypes, and the plaques in these mice were less compact and more diffuse in nature. Moreover, IF and scRNAseq analyses pointed to decreased numbers of PAM and DAM in CD33M^+^ microglia, which could account for less compaction. There are several possibilities on how CD33M may hinder DAM formation. Decreased DAM could be a secondary, long-term, consequence of impaired phagocytosis in CD33M^+^ microglia, which leads to elevated Aβ deposition. Another possibility is that CD33M directly impairs transcriptional reprogramming of microglia and CD33M is proposed to negatively regulate TREM2 [64, 65], which could be a mechanistic link. In this regard, CD33M microglia appear to be stalled at homeostatic (HM) and transitioning (TM) stages and fail to transition into DAM, which is reminiscent of mice lacking functional TREM2 [66]. A third possibility is that CD33M^+^ microglia initially differentiate into DAM but are driven into an alternative pro-inflammatory, detrimental, state due to overt activation and high proliferation [67]. Consistent with this idea, CD33M mice had increased IBA-1 levels and higher levels of Ki67^+^ in microglia. Higher IBA-1 levels are in line with a prior report of increase IBA-1 density from human AD patients homozygous for the AD-risk (rs12459419C) *CD33* allele [13]. In both our CD33M mice and this analysis of AD patients, higher IBA-1 density may be a secondary consequence of elevated Aβ levels, that leads to overly activated, yet less functional microglia. Indeed, constitutively activated microglia are detrimental and defective in the context of AD pathogenesis [67, 68].

We previously showed that the ability of CD33M to repress phagocytosis is dependent on its cytosolic ITIM [19], therefore, its role as a cell surface inhibitory receptor is likely a key contributing factor to the phenotypic outcomes we observe. It is interesting to speculate the potential role *trans* ligands of CD33M could have *in vivo.* This is especially worth exploring in the context of AD brain, as dramatic changes have been observed in the brain glycome of animal models of AD as well as human patients [69–71]. Recent advances in understanding human sialome and elucidation of the glycan ligands of CD33 [72–75] could be instrumental in spatial characterization of glycan ligands of CD33M at both regional and cellular levels in the brain, their changes during the course of disease, and their potential role in modulating microglia.

The decrease in total Aβ levels and increased plaque compaction in CD33m compared to control 5XFAD mice further support a gain-of-function role for CD33m *in vivo*. CD33m^+^ microglia had higher levels of internalized Aβ, which is in line with the ability of CD33m to enhance phagocytosis as we have previously reported [15]. Moreover, CD33m enhanced the number of PAM and increased microglial plaque encapsulation. Although scRNAseq analysis did not show dramatically enhanced differences in the number of DAM between CD33m and control mice, fewer Ab plaques in CD33m mice necessarily means a greater number of DAM per Ab cluster in CD33m. Indeed, higher density of Clec7a staining in PAM strongly suggest that CD33m had a higher number of DAM per Ab cluster.

Phagocytic clearance of small and intermediate Aβ assemblies can modify plaque composition in numerous ways such as discarding diffused Aβ that extend from the plaque core [41, 42, 76]. Therefore, enhanced phagocytic activity of CD33m^+^ microglia may be a key contributing factor to the AD protective effects of this isoform. However, there are no clear connections between enhanced phagocytosis and higher number of PAM/DAM, suggesting that the beneficial effects of CD33m may indeed be multifactorial. Proteomics, transcriptomics, and IF staining all consistently pointed to upregulated expression of nestin in CD33m^+^ microglia, and nestin was localized to microglia-plaque contact sites. Nestin is an intermediate filament abundantly expressed in cells with high migration capacity such as stem cells and is not normally expressed within homeostatic microglia. However, nestin has been shown to be upregulated in microglia under three circumstances: (i) following depletion of microglia when microglia are repopulating the brain [77], (ii) during inflammatory response in glial cells including activated microglia/macrophages [45], and (iii) in neonatal stages [78]. In common with all these conditions is cellular migration and, consistent with this concept, a previous study in cultured BV2 microglia expressing CD33m reported enhanced migration [16]. Beyond enhancing migration/retention of microglia to plaques, it cannot be ruled out that CD33m^+^ microglia also enhanced transcriptional reprogramming of microglia to a DAM phenotype. CD33m^+^ microglia from 5XFAD mice had upregulated IEGs in accordance with our previous observation under homeostatic conditions [15], although it is not obvious whether these relate to DAM formation. Concurrently, a functional role for CD33m could also be at play, as we have previously shown that the *in vitro* gain-of-function role for CD33m was dependent on its C-terminal cytosolic signaling motifs [15]. In this regard, it is interesting to speculate that the gain-of-function role of CD33m is governed by its unique, intracellular localization, reported by our group and others [15, 79].

Our results on neuronal health provide evidence that the impact of hCD33 isoforms on Aβ pathogenesis can go beyond total Aβ levels or plaque compaction. Specifically, changes in neuronal health were supported by: (i) differences in the size and number of dystrophic neurites; (ii) a GO signature associated with protected neural synaptic health in the mass spectrometry profiling; and (iii) behavior in the open field test. These effects are in line with many other studies pointing to the positive, neuroprotective impacts of an efficient microglial response upon neurodegeneration, especially at early stages of disease wherein microglia directly engage in clearance of neurotoxic Aβ deposits [80, 81]. Specifically, it has been shown that microglia limit plaque expansion by remodeling plaque morphology, through phagocytic trimming of diffused Aβ deposits that extend from the plaque core, making plaques more compact [41, 82]. Moreover, microglia envelop the plaque with their processes and form a physical barrier to block accumulation of neurotoxic, Aβ assemblies and stop their spread [40, 83]. Differences in plaque compaction have been shown to be associated with higher levels of Aβ induced neurotoxicity and worsened axonal pathology in humans as well as mouse models of amyloidosis [40, 41, 53, 84]. In this regard, investigating the impact of CD33 in a double pathology model that can recapitulate both the plaques and tangle pathologies, will be an important next step.

Interest in modulating microglial cell response as a therapeutic approach in AD has been gaining momentum [5, 85]. Specifically, approaches focusing on modification of identified susceptibility variants are of great interest. For example, agonistic anti-TREM2 antibodies have shown promising effects in pre-clinical mouse models [86–89]. Anti-CD33 antibodies targeting the extracellular domain of CD33M had also entered phase I clinical trials but stopped for unknown reasons. Although immunotherapy-based approaches can be considered useful for targeting AD risk variants, findings from our previous work and current study highlights the importance of considering both hCD33 protein isoforms in the context of therapeutic targeting of the *CD33* loci. In other words, an approach that only considers CD33M will miss the gain-of-function effect of CD33m, which are clearly stronger based on our observations. In this regard, it will be important to re-consider strategies that skew the formation of different hCD33 isoforms, such as splicing modulators [90]. A better understanding of the mechanistic aspect of CD33m gain-of-function is extremely crucial, as discovering the precise mechanism(s) will help open the door to new, promising therapeutic targets. An unexplored question is whether the gain-of-function role for CD33m converges on downstream effects of TREM2. In this regard, some of the transcriptional changes in CD33m mice, such as increased expression of several IEGs, resemble reported observations in mice treated with agonistic anti-TREM2 antibody [89]. Therefore, establishing whether enhancing CD33m expression converges on TREM2 and potentially drives synergistic effects with TREM2 agonism is a unique point to address.

The hCD33 Tg mouse models used in this study were generated with the aim to decipher the functional impact of individual isoforms in the context of Aβ pathogenesis. Although our findings clearly show a detrimental impact from CD33M and provides *in vivo* evidence towards protective, beneficial effects of CD33m it should be noted that the *CD33* gene, whether it is the rs12459419T or C allele, have both isoforms expressed. Using the same transgenic mice under homeostatic conditions, we previously showed that the gain-of-function role for CD33m in microglia is dominant over the suppressive effects of CD33M [15]. However, it is possible that the combination of the two isoforms could produce a phenotype in AD that is distinct from ones that can be generated for either isoform on its own. Additionally, while the focus of these studies was hCD33 isoforms, due to the significant differences between hCD33 and mCD33, we cannot fully rule out an effect from mCD33.

## Conclusions

Deciphering the functional aspect of AD susceptibility factors can lead to identification of pathways that can potentially be targeted for development of novel, promising therapeutics. The AD-protective effects of a minor *CD33* SNP motivated us to create a platform and independently investigate the loss-of-function and gain-of-function roles for the two CD33 protein isoforms associated with AD risk. Overall, our results suggest that AD susceptibility mediated by the *CD33* loci is a complex phenotype stemming from independent and opposite effects of hCD33 isoforms in modulating the microglial cell response to Aβ deposition. The divergent roles of hCD33 isoforms in microglia have an impact on total Aβ levels and plaque composition followed by downstream effects on neuronal health and cognitive function of the animals.

## Declarations

### Ethics approval

Ethics approvals are in place for all the animal work carried out in this study.

### Availability of data and materials

The RNA-seq expression data has been deposited to the GEO database. The full list of mass spectrometry data including peptides, and substrates identified is provided online (.xlsx) and also deposited in the MASSIVE repository.

### Authors’ contributions

G.E.S was involved in all aspect of this work including conceptualization, experimental design, experimental procedures, and data analysis. M.S.M. was involved in conceptualization, experimental design, and data analysis. M.C assisted with all stages of IF experiments including imaging and data analysis. S.Z and J.P performed the scRNAseq experiments. E.G and O.J performed the proteomics analysis. C.D.S assisted with sample preparation for cell sorting. L.M.C and V.S contributed to biochemical characterization of extracted Aβ assemblies. Z.P and Q.T performed the behavioral assays. G.S and V.A contributed to IF experiments. S.S assisted with mouse colony maintenance and genotyping. G.E.S and M.S.M wrote the manuscript and all authors read and approved the final manuscript.

## Acknowledgements

We would like to thank Dr. Mathew Blurton-Jones and Dr. Hayk Davtyan for helpful feedback and discussion. We thank Dr. Aja Rieger from the Faculty of Medicine and Dentistry Flow cytometry facility at University of Alberta for help with cell sorting. We also thank Dr. Jack Moore and Ramanaguru Siva for their help with mass spectrometry and proteomics data processing. We thank the staff and personnel of Health Sciences Laboratory Animal Services at University of Alberta for their help with animal maintenance.

## Abbreviations

AD: Alzheimer’s disease!
Siglec: sialic acid-binding immunoglobulin-type lectin
hCD33: Human CD33 protein
mCD33: murine/mouse CD33 protein
CD33M: Human CD33 major isoform
CD33m: Human CD33 minor isoform
GWAS: Genome-wide association studies
SNP: Single nucleotide polymorphism
Siglec: Sialic acid-binding immunoglobulin-type lectin
Aβ: Amyloid beta
ITIM: Immunoreceptor tyrosine-based inhibitory motif
ITAM: Immunoreceptor tyrosine-based activatory motif
IF: Immunofluorescence microscopy
SCCAF: Single-Cell Clustering Assignment Framework
pySCENIC: Python Single-Cell rEgulatory Network Inference and Clustering
UMAP: Uniform manifold approximation and projection
IEG: Immediate early gene
scRNAseq: Single cell RNA sequence
DEGs: Differentially expressed genes
DAM: Disease associated microglia
DIA-MS: Data-independent acquisition mass spectrometry
PAM: Plaque associated microglia
BAM: Border associated macrophages
TM: Transitioning microglia
RBM: RNA binding protein microglia
HM: Homeostatic microglia
GO: Gene ontology

