## Supplementary material for "Alzheimer’s disease associated isoforms of human CD33 distinctively modulate microglial cell responses in 5XFAD mice": Suppl Figures

### Suppl. Figure 1

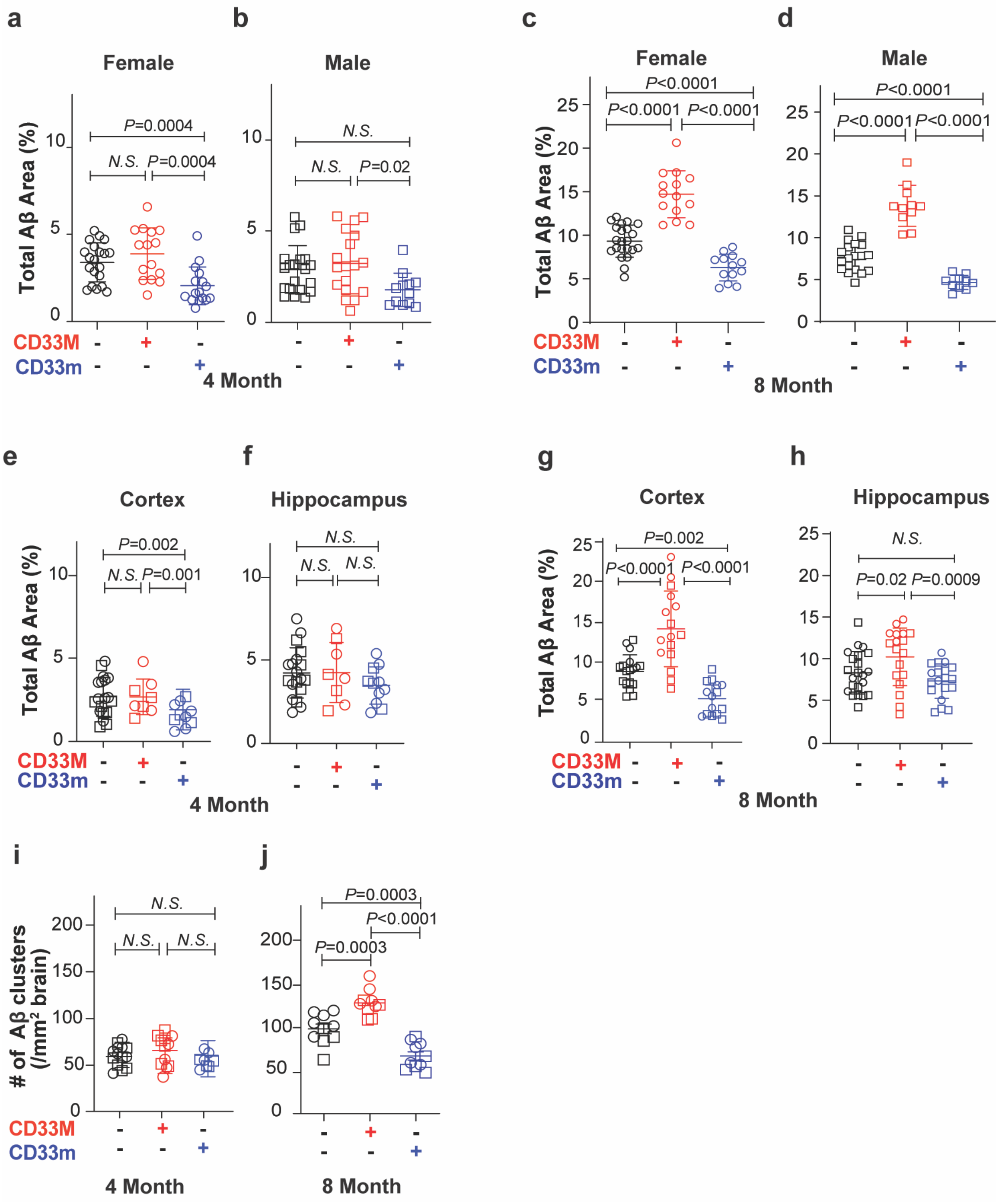

### Suppl. Figure 2

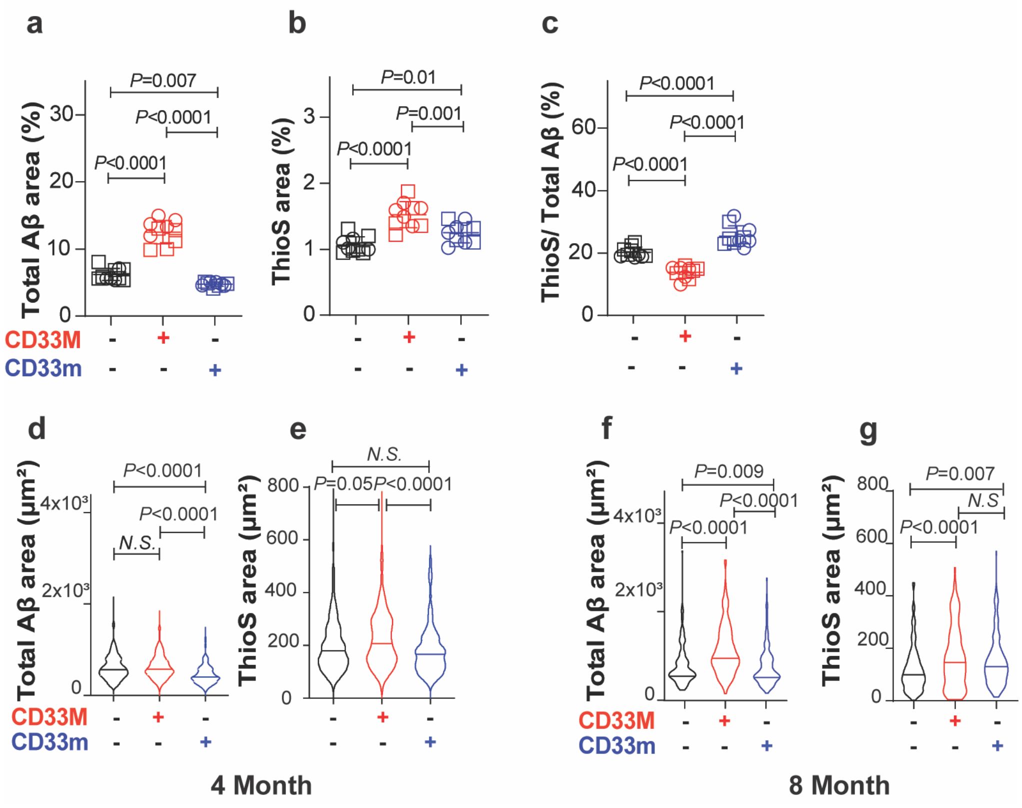

### Suppl. Figure 3

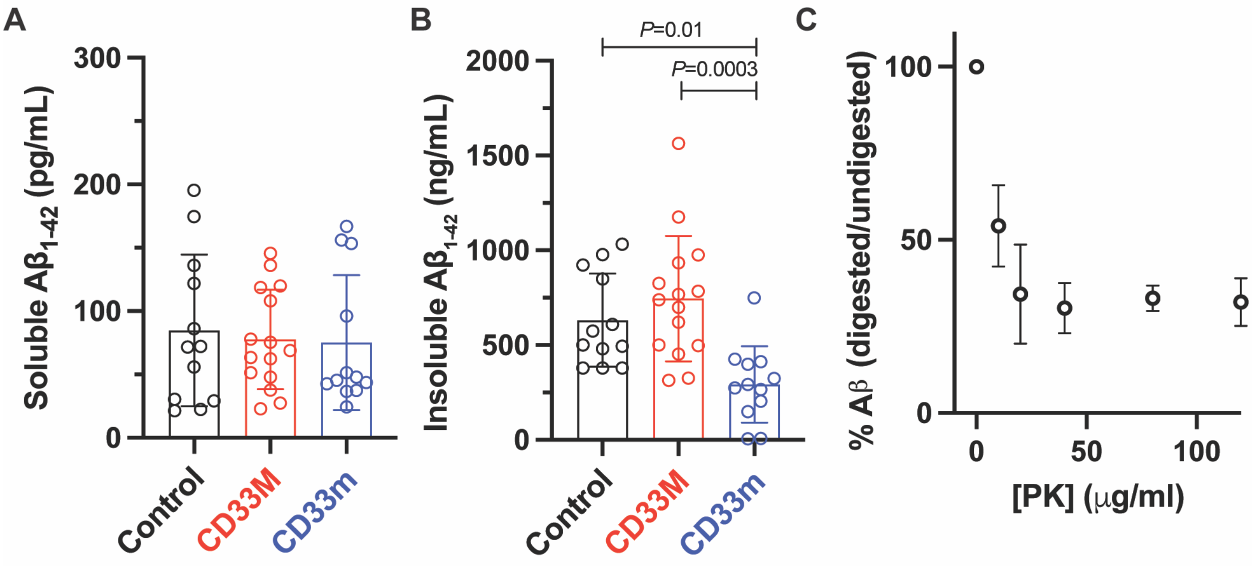

### Suppl. Figure 4

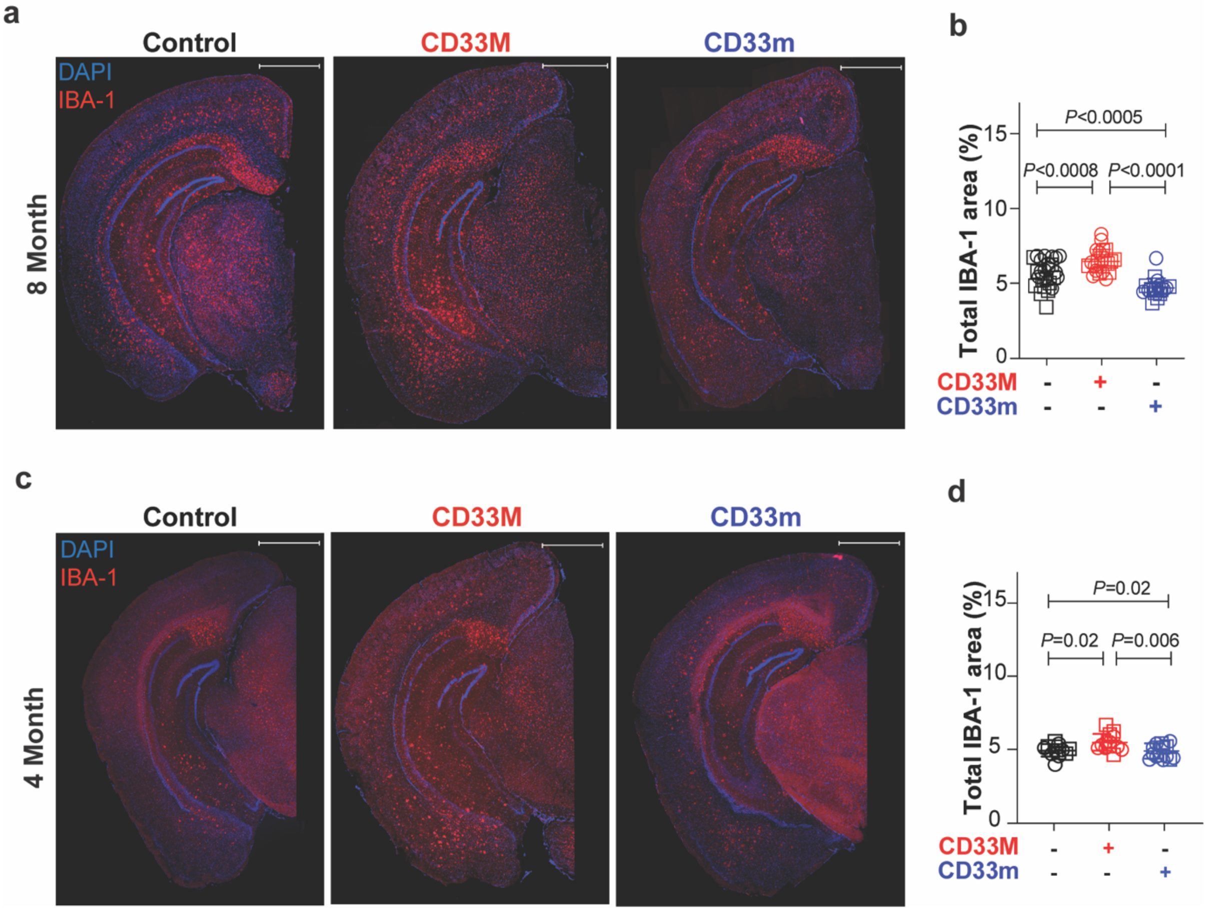

### Suppl. Figure 5

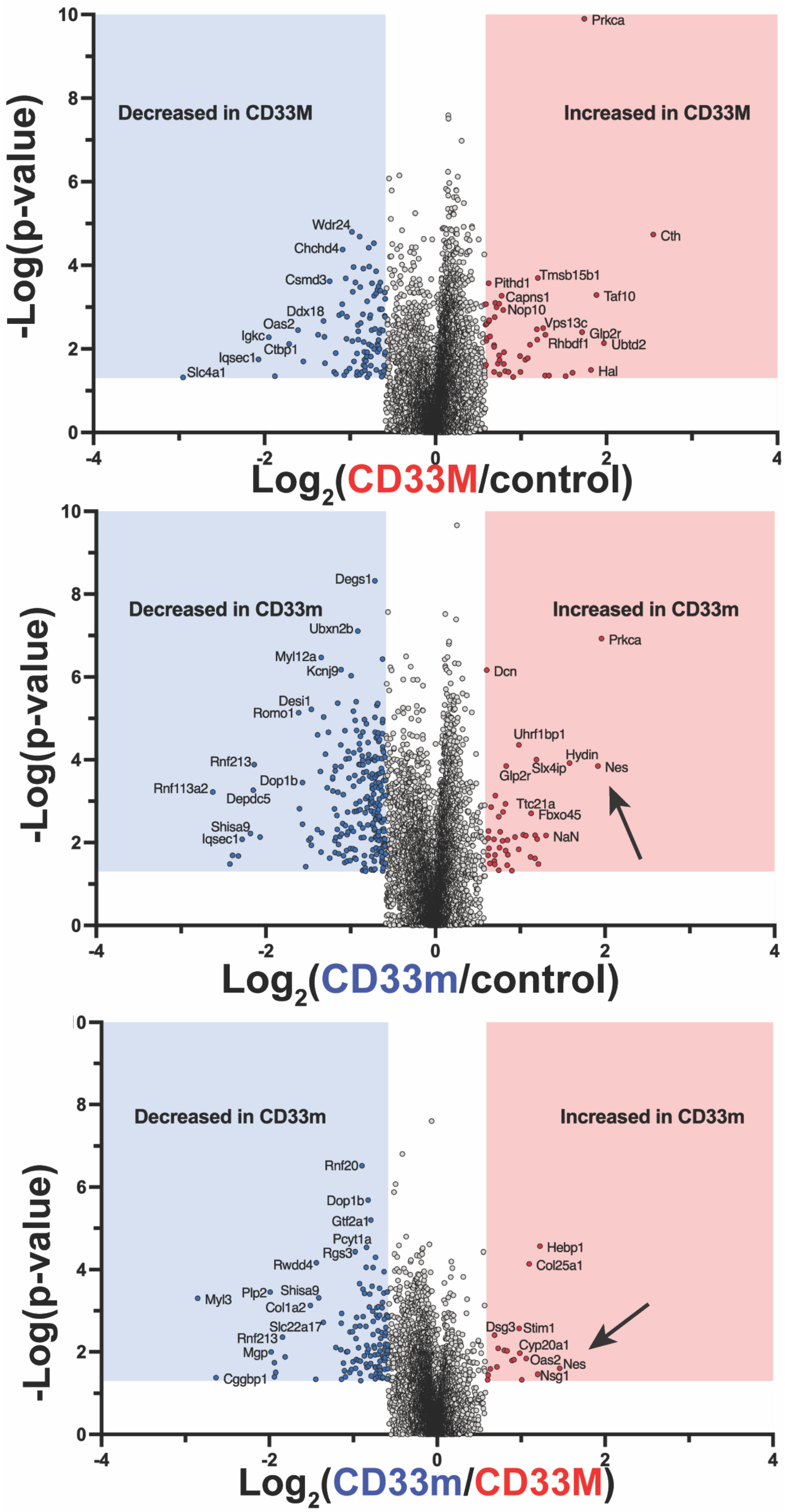

#### Suppl. Figure 6

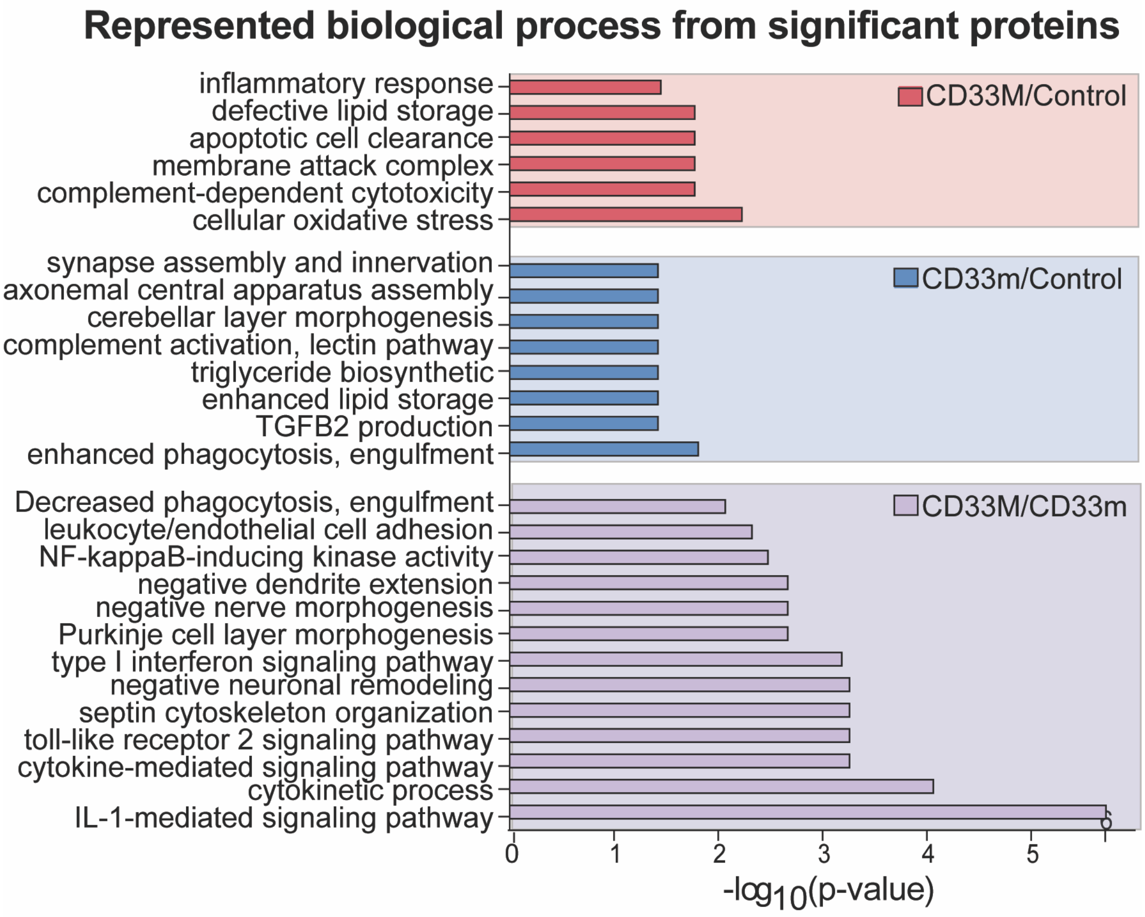

### Suppl. Figure 7

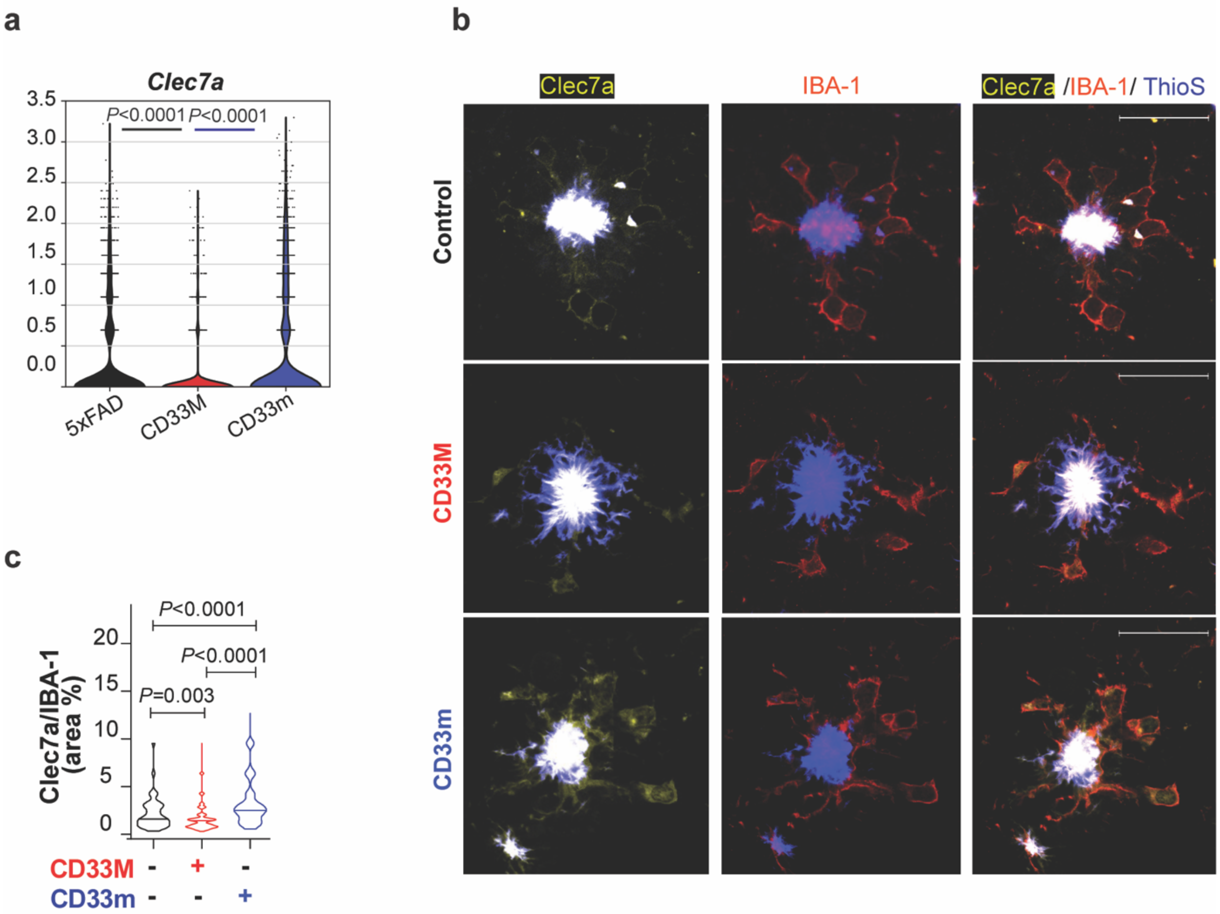

### Suppl. Figure 8

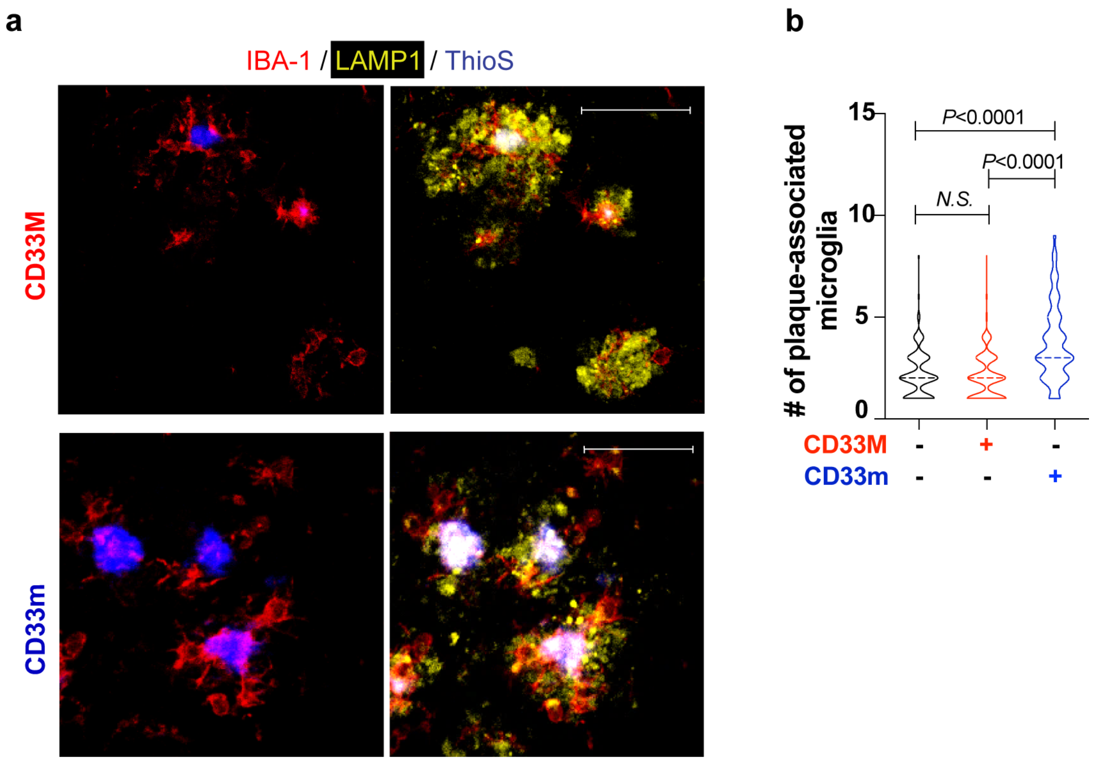
